# Proteogenomic Characterization Reveals Therapeutic Opportunities Related to Mitochondrial Function in Melanoma

**DOI:** 10.1101/2022.10.24.513481

**Authors:** Jeovanis Gil, Yonghyo Kim, Viktória Doma, Uğur Çakır, Magdalena Kuras, Lazaro Hiram Betancourt, Indira Pla Parada, Aniel Sanchez, Yutaka Sugihara, Roger Appelqvist, Henriett Oskolas, Boram Lee, Jéssica de Siqueira Guedes, Gustavo Monnerat, Gabriel Reis Alves Carneiro, Fábio CS Nogueira, Gilberto B. Domont, Johan Malm, Bo Baldetorp, Elisabet Wieslander, István Balázs Németh, A. Marcell Szász, Ho Jeong Kwon, Runyu Hong, Krzysztof Pawłowski, Melinda Rezeli, József Tímár, David Fenyö, Sarolta Kárpáti, György Marko-Varga

## Abstract

The dynamics of more than 1900 mitochondrial proteins was explored through quantitative proteomics in 151 melanoma-related tissue samples of both surgical and autopsy origin. Dysregulation of mitochondrial pathways in primary tumors, metastases, and peritumoral tissues was correlated with age and survival of patients, as well as with tumor cell proliferation and the BRAF mutation status of the tumors. The outlined proteomic landscape confirmed the central role of a pathologically upregulated mitochondrial translation machinery and oxidative phosphorylation (OXPHOS) in the development, proliferation, and progression of melanomas. Our results from different melanoma cell lines confirmed our findings and we could document that treatments with selected OXPHOS inhibitors and antibiotics successfully impaired tumor cell proliferation. In addition, we provided proteomic evidence on the mechanism-of-action of the different treatments. These observations could contribute to the development of therapeutic approaches targeting the mitochondrial pathology in melanoma.

**TOC figure:** 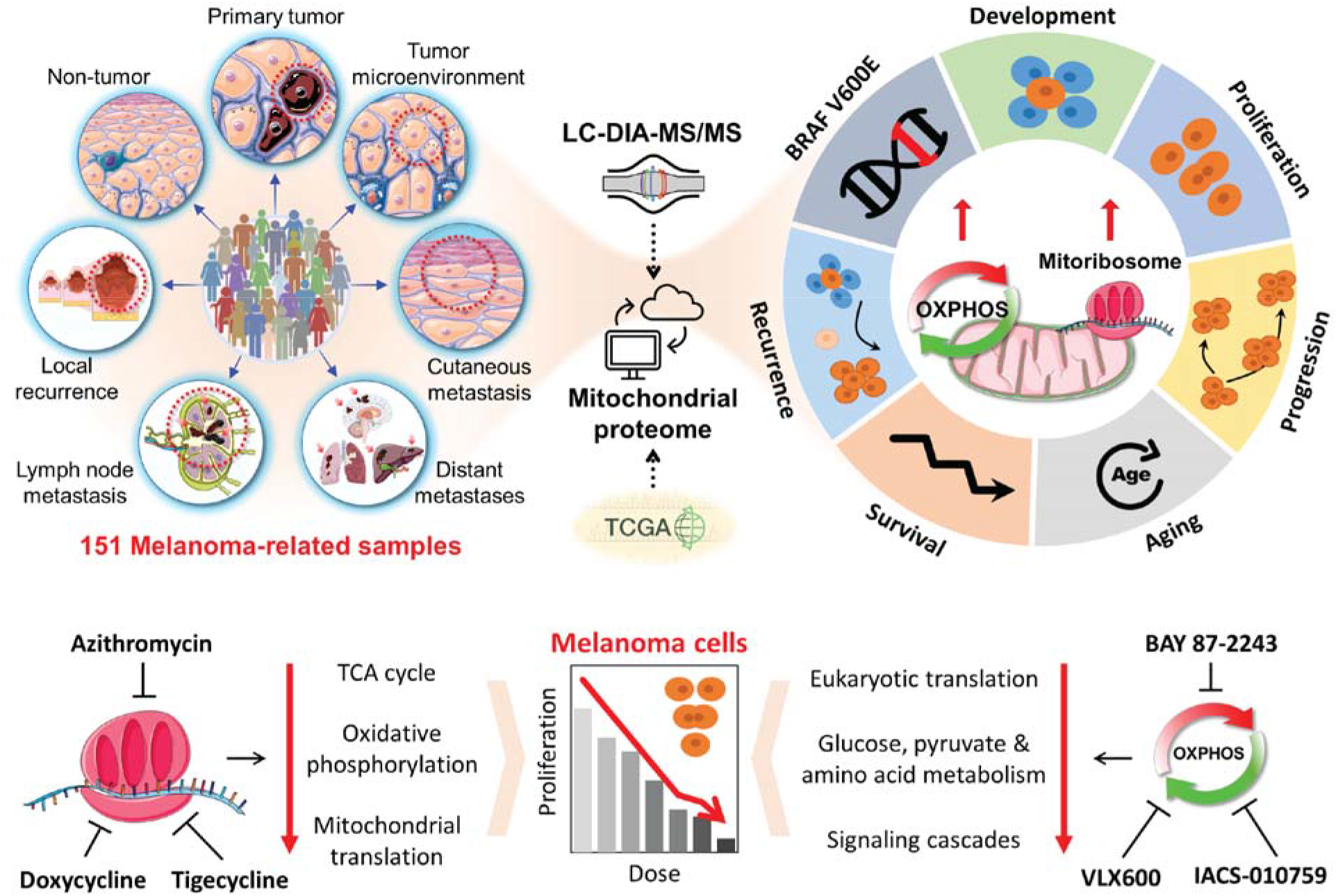

**Highlights:**

- Mitochondrial proteome landscape outlined in 151 melanoma-related samples
- Mitochondrial Translation and OXPHOS impact disease severity and survival
- BRAF V600E mutation correlates with upregulation of mitochondrial energy production
- Targeting the mitochondrial OXPHOS and ribosomes impairs tumor cell proliferation
- Therapeutic opportunities complementary to the standard of care are proposed

**In brief:** Mitochondrial proteome profiling of melanomas reveals dysregulation in major metabolic pathways, suggesting a central role of the mitochondria within the development and progression of melanoma. Targeting mitochondrial pathways has the potential to impact the course of the disease, which provides opportunities for complementary drug interventions.

## 1. Introduction

Melanoma is considered the most aggressive skin cancer of relatively frequent occurrence. It develops from pigment-containing cells known as melanocytes (Dimitriou et al., 2018). While the most important factor in cutaneous melanoma development is still the skin exposure to environmental UV light, the risk increases when combined with low levels of skin pigment, large numbers of pigmented nevi, other genetic and environmental factors, and/or with a compromised immune system, which can all contribute to tumorigenesis, and in rare cases, also extracutaneously (Erdei and Torres, 2010; Kaiser et al., 2020). According to the GLOBOCAN database, the estimated worldwide incidence of melanoma falls in the 19^th^ place among cancers, while in Europe, the ranking rises to the 6^th^ position (Ferlay et al., 2020; Leonardi et al., 2018). In 2020 of all diagnosed cancers in Europe, 3.7% were melanomas. In addition, about 17.5% of all diagnosed melanomas lethally relapsed following treatment. The main prognostic factor for melanoma, the histopathological classification and staging, has primarily relied on the tumor thickness and the presence of the sentinel lymph node status, and it is a major estimate of the clinical behavior of primary melanomas (Gershenwald et al., 2017). While a melanoma gene expression profile (MGEP) has been designed to predict prognosis from the primary tumor, it is not incorporated yet into the clinical practice (Grossman et al., 2020). The main driver gene mutations to date have been identified in the BRAF, NRAS, KIT, NF1, GNAQ, TP53, TERT, and PTEN genes. These markers, however, cannot act as individual indicators for every melanoma case. Our recent research highlighted that, in addition to the mutation status at the gene level, the actual amount of mutated protein has a critical impact on tumor biology and ultimately on patient survival (Betancourt et al., 2019). Furthermore, melanoma is one of the cancers with high genetic heterogeneity which results in high variability in the disease phenotypes (Alexandrov et al., 2013). This high level of heterogeneity represents a key element for the progression of the disease and has been linked to shorter survival of patients (Gara et al., 2018; Szász et al., 2017).

The metabolic shift from oxidative phosphorylation (OXPHOS) to a glycolytic phenotype has been considered a hallmark of cancer (Hanahan and Weinberg, 2011). However, there is also an increasing number of studies linking mitochondrial pathways, particularly OXPHOS, to cancer development and progression (Danhier et al., 2017; Porporato et al., 2018; Roberts and Thomas, 2013). Mitochondrial metabolism has been reported to impact cancer development in several ways. The increased generation of Reactive Oxygen Species (ROS) as a byproduct of the OXPHOS can support the genomic instability required for malignant transformation. Intermediate metabolites from mitochondrial metabolism are required for rapid cell proliferation, including some with intrinsic transforming properties (Porporato et al., 2018). Particularly in melanoma, the resistance to BRAF target therapy was associated with a metabolic switch towards an OXPHOS phenotype to adapt their metabolism (Trotta et al., 2017). Interestingly, a large proteomic study revealed that tumors from patients responders to immunotherapy upregulated the OXPHOS and lipid metabolism (Harel et al., 2019). More recently, a small study on primary tumors revealed a progression dependence on the upregulation of mitochondrial ribosomes and oxidative phosphorylation (Gil et al., 2021). Thus, targeting mitochondrial functions is gaining attention in cancer treatment. The anticancer properties of OXPHOS inhibitors have been recently evaluated in several cancer models including melanoma, acute myeloid leukemia, brain and colon cancer (Gopal et al., 2019; Molina et al., 2018; Zhang et al., 2014). The mitochondrial translation machinery is responsible for the biosynthesis of 13 OXPHOS proteins encoded in the mitochondrial genome. Mitoribosomes are composed of two subunits that show homology to their bacterial ancestors (Zimorski et al., 2014). Moreover, several antibiotics approved for targeting bacterial protein synthesis also inhibit mitochondrial translation. Recently, this class of antibiotics has been used to test the functional dependence of cancer cells on mitochondrial translation. Targeting mitoribosomes has shown high selectivity in inhibiting the proliferation of cancerous cells in many cancer types including leukemias and solid tumor models (Kim et al., 2017; Lamb et al., 2015; Škrtić et al., 2011).

The present study aimed to analyze the mitochondrial proteome dynamics in 151 melanoma-related tissue samples. Mitochondrial proteome was filtered from identified proteins based on their evidence in MitoMiner repositories. Dysregulations in mitochondrial pathways were associated with melanoma development, proliferation of tumor cells, the progression of the disease after treatment, the age and survival of the patients, and with the BRAF mutation status of the tumors. Particularly, upregulation of mitochondrial OXPHOS and translation machinery was evident in the more aggressive presentation of the disease, which strongly suggests that the mitochondria play a driving role in the development and progression of melanoma. The major findings were confirmed at the transcriptomic level in a separate dataset of 443 melanomas publicly available on the TCGA repositories. The results from targeting both the mitoribosomes and the OXPHOS in melanoma cell culture models showed a drug dose-dependent impairment of the proliferation of melanocytes. Furthermore, in-depth proteomics revealed details on the molecular mechanism-of-action of the tested drugs.

## 2. Results

Quantitative proteomic analyses were performed on mitochondrial proteins from 151 melanoma-related samples. A total of 77 surgically resected tissue samples were obtained from 47 melanoma patients (prospective cohort). The samples were classified based on their origin; non-tumor (NT; >2 cm from the tumor edges; n=11), tumor microenvironment (TM; tumor edge without microscopic evidence of tumor cells; n=6), primary tumor (PT; n=16), local recurrences (LR; n=7), cutaneous metastases (CM; n=9), regional lymph node metastases (LN; n=23), and distant metastases (DM; coming from gallbladder, brain, liver, spleen, and breast; n=5) (**Figure 1A**). Additional 74 tissue samples were life-threatening metastases in 20 different organs from 22 deceased melanoma patients (postmortem cohort). The number of metastases per patient ranged from 1 to 9, lung (n=14) and liver (n=11) being the most commonly affected organs (Doma et al., 2019) (**Figure 1A**). All samples were snap-frozen within 120 minutes of harvesting and stored in a fully integrated and automated large-scale biobank (Fehniger et al., 2013; Malm et al., 2015, 2018; Marko-Varga et al., 2012).

**Figure 1.**
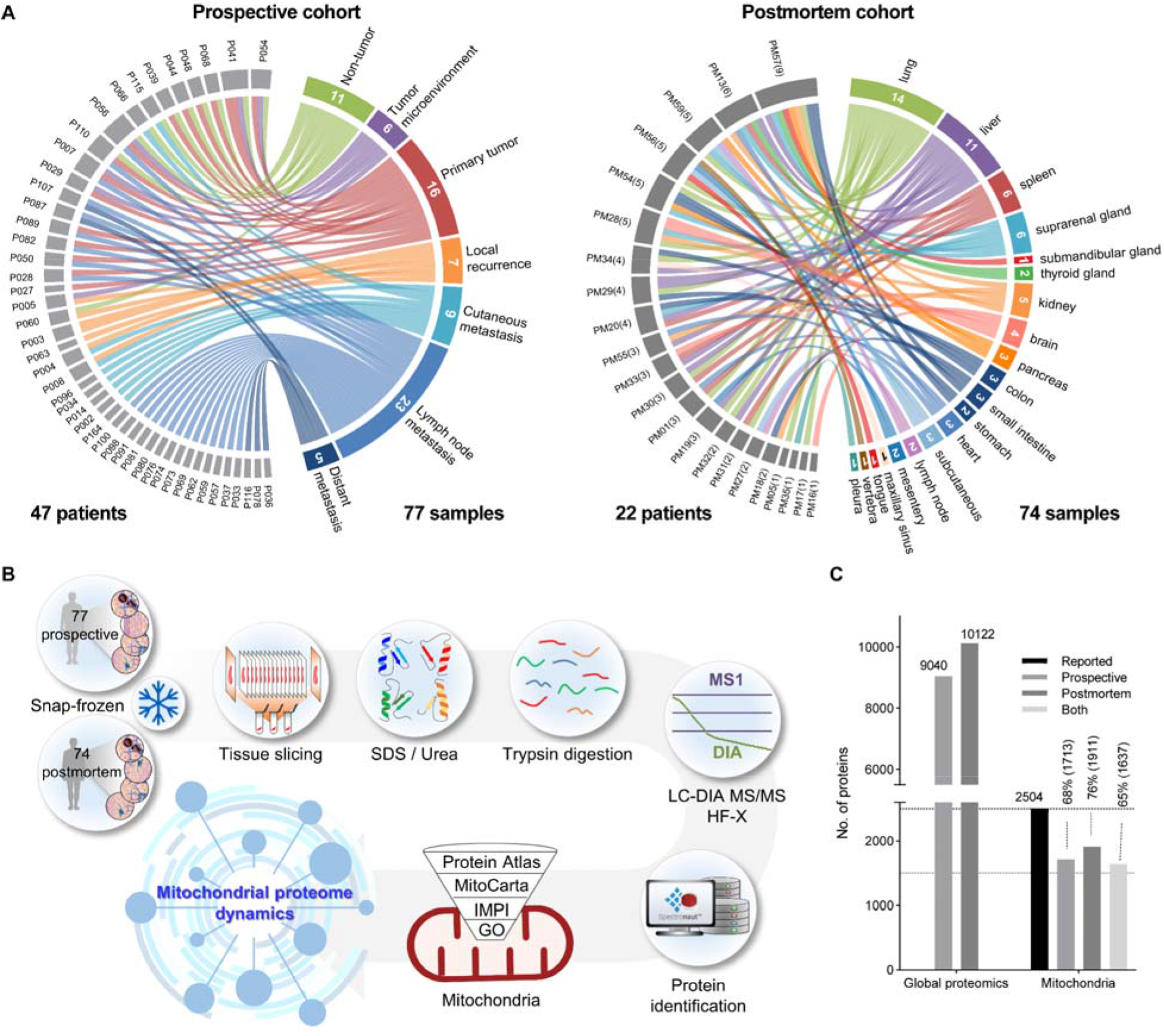
Melanoma study outline. **A)** Distribution of the samples included in the prospective (left) and the postmortem (right) cohorts. The samples in the prospective cohort originated from non-tumor, tumor microenvironment, primary tumor, local recurrence, cutaneous, lymph node, and distant metastases and were obtained from 47 patients. The samples included in the postmortem cohort; all were melanoma metastases in 20 different organs from 22 different individuals. Connections link samples with patients. **B)** Sample preparation strategy and proteomic workflow followed by filtering the mitochondrial proteome and bioinformatic analysis. **C)** Total numbers of proteins identified in the study and those reported to be mitochondrial according to MitoMiner.

Samples were processed according to previous reports (Gil et al., 2017a, 2019; Kuras et al., 2019) and analyzed with high-resolution mass spectrometers (QExactive HF-X) using a Data Independent Acquisition (DIA) method (**Figure 1B**). We confidently identified and quantified 9040 and 10122 proteins from the prospective and postmortem cohorts respectively, which provided the basis for performing deep-mining into the disease biology. Mitochondrial proteins were filtered by querying the MitoMiner 4.0 database, which takes evidence from four major repositories to report the mitochondrial location of proteins: MitoCarta, Integrated Mitochondrial Protein Index (IMPI), Gene Ontology Resource (GO), and the Human Protein Atlas (Calvo et al., 2016; Pagliarini et al., 2008), respectively. Of the 2504 proteins reported as mitochondrial, 1713 and 1911 were identified in the prospective and postmortem cohorts respectively, including 1637 common to both datasets (**Figure 1C**). Further analyses were performed with the standardized quantitative data of the mitochondrial proteome.

### 2.1. Mitochondrial metabolic reprogramming in melanoma

By analyzing the mitochondrial protein profiles in the prospective cohort, a total of 651 proteins showed significant differences based on their sample origin (ANOVA, FDR<0.05). Subsequent **post hoc** t-test analyses revealed that most of these proteins were indeed dysregulated between the non-tumor group and the groups of tumors, and to a lesser extent, the tumor microenvironment samples. Differential abundance profiles showed, in all cases, an excess of proteins significantly upregulated relative to the downregulated by comparing the tumor- and microenvironment-with non-tumor groups (**Figure 2A**). A total of 237 proteins were significantly upregulated in all tumors compared to non-tumor samples. Furthermore, 228 of these also showed higher levels in the tumor microenvironment, including 79 significantly upregulated, in comparison to the non-tumor samples. These findings indicate a profound change in the mitochondrial proteome during melanoma development, extending beyond tumor cells into the microenvironment.

**Figure 2.**
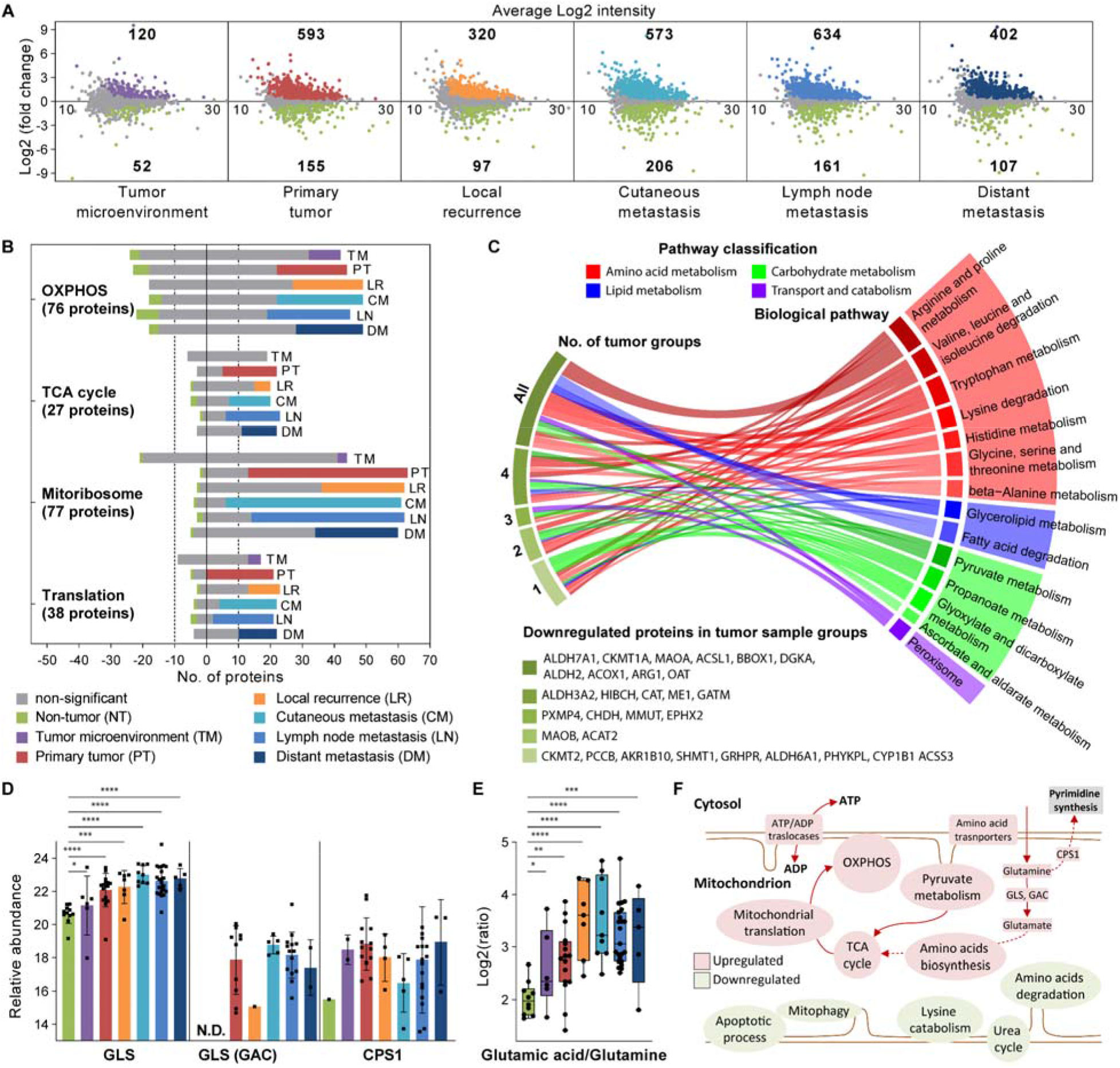
OXPHOS, TCA cycle pathways, and the mitochondrial translational machinery are upregulated, and amino acid and lipid metabolism are downregulated in melanoma tumor samples compared to non-tumors. **A)** Median average (MA)-plots showing mean differences in mitochondrial protein abundances between non-tumor and all other groups; significantly up- and downregulated proteins defined as FDR q value <0.05 are plotted in colors other than gray and the numbers are represented, negative log2 fold change values indicate downregulated proteins compared to non-tumor samples. **B)** Distribution of OXPHOS proteins, the TCA cycle, and mitochondrial translational machinery proteins, considering their relative level compared to the non-tumor group. Negative and positive bars indicate the number of downregulated and upregulated proteins in the microenvironment or tumor groups. Colors other than gray refer to significantly dysregulated proteins. **C)** Significantly enriched pathways in proteins downregulated in at least one tumor group (Primary, Local recurrence, Cutaneous, Lymph node, and Distant metastases) relative to non-tumor. Shades of green indicate in how many groups of tumor samples the represented proteins are downregulated. **D)** Protein abundance profiles in the different groups of the mitochondrial glutaminase kidney isoform (GLS) and its isoform 3 (GAC) and the mitochondrial carbamoyl-phosphate synthase [ammonia] (CPS1). **E)** Abundance ratio of glutamic acid and glutamine in the tissue samples. Significant differences between the non-tumor group and all others were estimated through one-way ANOVA and the cut-off was set based on a FDR approach, (*,**,***,****) q value < 0.05, 0.01, 0.001, 0.0001 respectively). **F)** Schematic representation of the mitochondrial pathways dysregulated in melanoma.

The proteins commonly upregulated in both the tumor and microenvironment mainly belong to pathways such as OXPHOS, TCA cycle, and the mitochondrial translational machinery. The protein profiles of these pathways showed that the vast majority were upregulated in all groups compared to non-tumor, although not all were statistically significant (**Figure 2B**). These findings indicated that melanoma cells and their microenvironment depend on mitochondrial pathways to ensure their energy requirements, in addition to the upregulation of the glycolysis, as previously reported (Gil et al., 2019). SLC25A4, SLC25A5 and SLC25A6 out of the four known ADP/ATP translocases responsible for the exchange of cytosolic ADP with mitochondrial ATP showed relatively higher expression within the tumor groups, which also supports our hypothesis (**Figure S1**). Interestingly, only peptidyl-tRNA hydrolase ICT1, which dictates the mitochondrial translation termination (Akabane et al., 2014), was significantly downregulated in all groups compared to non-tumors. This finding suggests that melanoma, unlike other cancers with ICT1 upregulation (Lao et al., 2016; Wang et al., 2017; Xie et al., 2017), is dependent on the other mitochondrial class 1 release factors MRTF1, MTRF1L, and MTRFR, which were only detected within the microenvironment and tumor samples (**Figure S2**).

Mitochondria are dynamic organelles that constantly undergo two processes: fission and fusion, in response to cellular requirements. Our results indicate an upregulation of the mitochondrial division process (fission) in melanomas. The fission regulation proteins including dynamin-1-like protein (DNM1L), mitochondrial fission factor (MFF), mitochondrial fission protein 1 (FIS1), and mitochondrial dynamics protein MID51 (MID51), were significantly upregulated in tumor groups (**Figure S3A**). This is supported by the increase of the E3 ubiquitin-protein ligase MARCH5 expression, which plays a critical role in controlling the mitochondrial morphology by positively regulating mitochondrial fission, and preventing cell senescence (Park et al., 2010; Xu et al., 2016).

We identified mitofusin-1 and -2 (MFN1 & MFN2), dynamin-like 120kDa protein (OPA1), and optic atrophy 3 protein (OPA3), which are key drivers of the mitochondrial fusion. However, only OPA1 and OPA3 were found to be upregulated in most of the tumor groups compared to non-tumors (**Figure S3B**). Besides, two mitochondrial sirtuins, SIRT3 and SIRT5, were upregulated in the tumor groups compared to non-tumors (**Figure S3C**). Mitochondrial sirtuins remove the damaging acetylation marks of the mitochondrial proteins and thus contribute to the preservation of the mitochondrial function (Gil et al., 2017b). Consistently with our findings, the SIRT5 gene was found with amplification/gain in 55% of melanomas according to the genomic data publicly available at the TCGA repositories. Moreover, silencing SIRT5 induces apoptosis in melanoma cells and inhibits tumor growth in xenografts (Giblin et al., 2021).

### 2.2. The metabolism of amino acids is dysregulated in melanoma

A total of 253 proteins were significantly downregulated in at least one tumor group compared to non-tumors. Most of these proteins are involved in amino acids, lipids, and to a lesser extent, carbohydrates metabolism pathways (**Figure 2C**). However, 37 of them are linked to apoptotic processes including p53 protein and some downstream effectors such as VDAC1, which functions as a releaser of mitochondrial products that triggers apoptosis (Li et al., 2014b). Downregulation of VDAC1 activates AMPK, resulting in mTOR suppression and autophagy induction that may contribute to the survival of tumor cells (Head et al., 2015; Hwang et al., 2020).

The two amine oxidases; MAOA and MAOB were downregulated in tumor samples. MAOA has been studied extensively in neurosciences, but very few studies have been conducted in cancer research and the results were different depending on the cancer type. In prostate cancer, its expression is strongly associated with disease progression and short survival, whereas the opposite was observed in hepatocellular carcinoma and cholangiocarcinoma (Flamand et al., 2010; Huang et al., 2012; Li et al., 2014a; Pang et al., 2020; Wu et al., 2014). Unlike melanoma, MAOB was found to be more active in gliomas (Gabilondo et al., 2008) and has recently been associated with poor prognosis in colorectal cancer (Yang et al., 2020). Furthermore, other proteins also involved in amino acid metabolism were downregulated in melanomas such as choline dehydrogenase (CHDH), alpha-aminoadipic semialdehyde dehydrogenase (ALDH7A1), and the two mitochondrial creatine kinases (CKMT1A and CKMT2). CHDH plays a role in mitophagy and its downregulation in melanomas might prevent mitochondrial degradation through autophagy as recently reported (Park et al., 2014). The knock-down of the cytosolic creatine kinase CKB has been reported to enhance mitochondrial activity (Li et al., 2013); thus, we might suggest a similar function for the downregulation of CKMT1A and CKMT2.

The presented data point toward a higher consumption of glutamine through its conversion to glutamate by melanomas. The glutaminase kidney isoform (GLS) was upregulated in all tumor groups compared to non-tumors. Its splice variant glutaminase C (GAC), which has a greater affinity toward glutamine (Cassago et al., 2012; Møller et al., 2013), was detected in tumor samples only (**Figure 2D**). Glutamine is the most abundant amino acid in human blood and is known to be highly consumed by tumor cells (Choi and Coloff, 2019). The levels of free amino acids and metabolites were also determined in the samples, where higher levels of glutamine were detected in most groups compared to non-tumors; however, the increase was significant only in primary tumors (**Figure supplemental 2E**). The levels of glutamate **(Figure S4)** and more importantly, the ratio between glutamate and glutamine were significantly higher in all tumor groups, as well as in the microenvironment (**Figure 2E**). Additionally, cysteine, phenylalanine, and leucine/isoleucine were found to be significantly more abundant in some tumor groups compared to non-tumors. On the other hand, arginine, serine, and histidine were less abundant in tumor samples (**Figure S4**). Mitochondrial carbamoyl-phosphate synthase (CPS1), which converts ammonia into carbamoyl-phosphate, was detected in most of the tumor samples while in only one of the non-tumors (**Figure 2D**). CPS1 is known to be affected by lysine acetylation and its deacetylation by SIRT5 activates the protein, contributing to the detoxification of ammonia (Hallows et al., 2009; Nakagawa et al., 2009). Carbamoyl phosphate is an important precursor for the synthesis of pyrimidines required for the proliferation of cancer cells and a source of ammonia to enter the urea cycle (Keshet et al., 2018).

Conclusively, this suggests that the relatively lower levels of the urea cycle enzymes in melanomas influence the downregulation of the pathway. Proteins such as arginase-1 (ARG1), which produces urea and ornithine from arginine, and argininosuccinate synthase (ASS1), which converts citrulline and aspartate to argininosuccinate, as well as ornithine aminotransferase (OAT), which transfers an amino group from ornithine to 2-oxocarboxylates, were downregulated in all tumor groups (**Figure S5A**). Additionally, the levels of ornithine were slightly lower in tumor groups and were only significant in lymph node metastases, while citrulline and citrulline/ornithine ratios were significantly lower in all groups compared to non-tumors (**Figure S5B-D**). Taken together, these findings suggest that the carbamoyl phosphate production in tumors is primarily directed towards proliferation-related pathways. Melanoma dysregulates mitochondrial pathways to support its development and progression, redirecting the metabolism to support the OXPHOS and the production of intermediates of proliferation **(Figure 2F)**.

Melanomas rely on the oxidation of glutathione to keep their production of reactive oxygen species (ROS) in balance. Upregulation of two members of the glutathione S transferase family (GSTO1 and GSTK1) together with glutathione peroxidase (GPX1) was observed in the study (**Figure S7A**). Interestingly, the levels of reduced glutathione (GSH) did not show significant differences between the groups, while on the other hand, oxidized glutathione (GSSG) levels were significantly higher and their ratios (GSH: GSSG) were significantly lower in all groups compared to non-tumor samples (**Figure S7B**). Taken together, these results suggest that the excess of ROS as byproduct from the upregulation of mitochondrial OXPHOS in melanomas and their microenvironment would be at least quenched through the oxidation of GSH. A previous report showed that by targeting two of the glutathione S transferase family members (GSTP1 and GSTM3), cancer cell proliferation and tumor size were inhibited in a xenograft model of cervical cancer (Checa-Rojas et al., 2018). The upregulation of GSTO1 and GSTK1 in melanomas might prevent apoptotic signaling triggered by the excessive ROS production from mitochondrial OXPHOS. These findings, with melanoma development dependence on the glutathione metabolism, may open a novel area of therapeutic opportunities.

### 2.3. Melanoma proliferation relies on mitochondrial translation

The minichromosome maintenance (MCM) protein complex is composed of six elements MCM2-7 and is required for DNA replication. The six proteins form a heterohexameric complex with an 1:1:1:1:1:1 stoichiometry, which has DNA helicase activity (Das et al., 2014; Lee and Hurwitz, 2001; Miller et al., 2019; You et al., 1999). Independently, all these proteins have been reported to be upregulated in several cancers and associated with fast progression, proliferation, aggressiveness and short survival (Aporowicz et al., 2019; Cao et al., 2020; Das et al., 2014; Giaginis et al., 2009, 2011; Issac et al., 2019; Kang et al., 2022; Lee and Hurwitz, 2001; Stojkovic-Filipovic et al., 2016; Valverde et al., 2018; Wojnar et al., 2010; You et al., 1999). The relative abundances, extracted from MS data of the six elements of the MCM complex, were taken as a measurement of the proliferation rate of the cells in all samples included within the study. A hierarchical clustering analysis of the samples using the protein abundances of the MCM complex was performed and four groups were created assuming that higher levels correlate with higher proliferation rate. The groups created were quantitatively ranked from Low-, to High-, including Medium-Low-, and Medium-High-proliferation. The resultant clustering analyses indicated that tumor samples experienced different proliferation rates independently from the sample origin (**Figure 3A**). However, as expected non-tumor and tumor microenvironment samples from the prospective cohort, did all fit into the Low and Medium-Low proliferative groups. Additional analysis was conducted on the RNA sequencing data from 443 melanomas publicly available at the TCGA repositories (Akbani et al., 2015). The proliferation clusters were created based on the transcript levels of the MCM complex elements (**Figure 3A**). In the RNA sequencing data, 2286 transcripts coding for mitochondrial proteins were identified.

**Figure 3.**
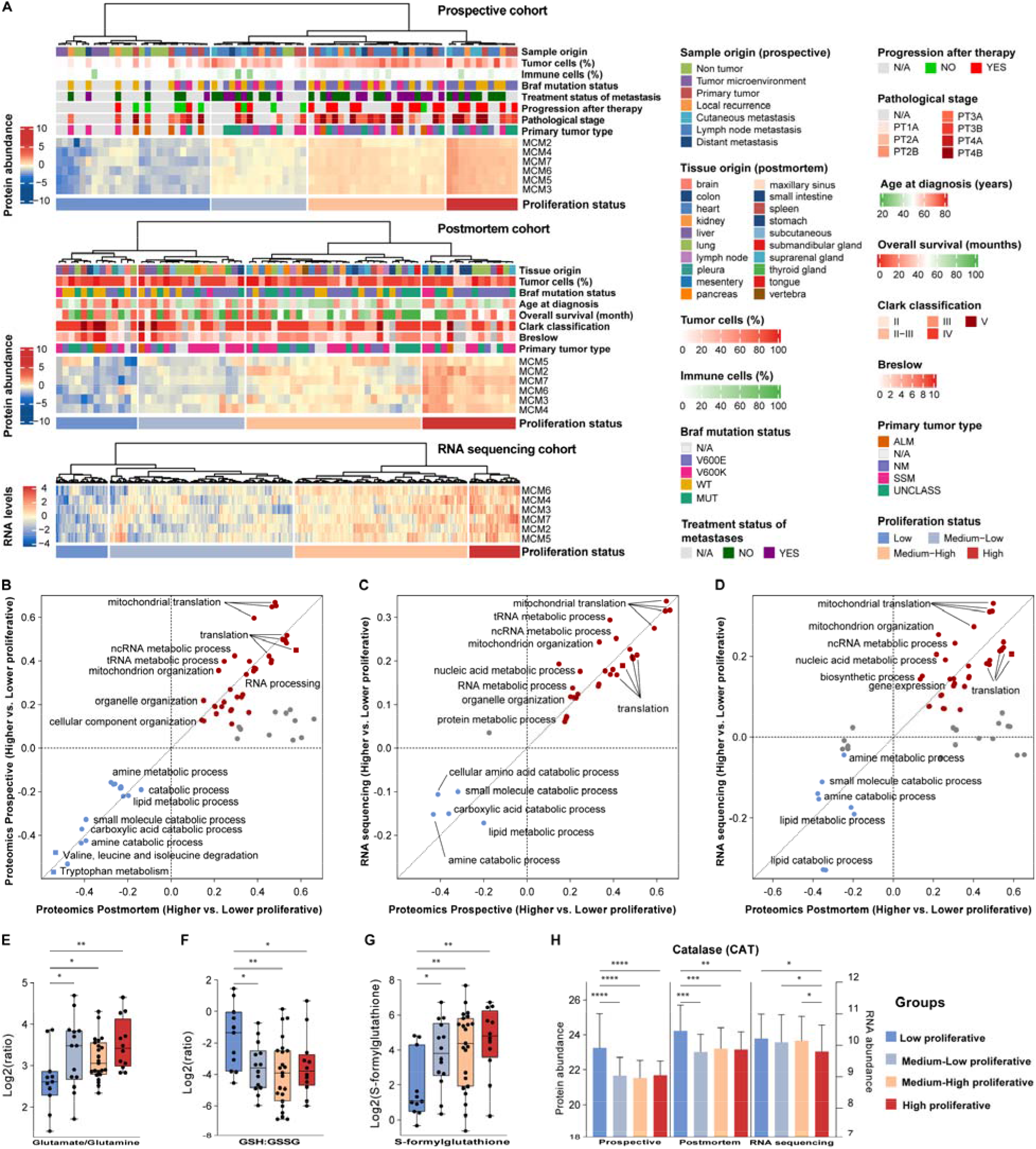
Tumor proliferation positively correlates with the mitochondrial translation machinery in melanoma. **A)** Unsupervised hierarchical clustering of tumor samples based on the relative expression abundance of the six elements of the MCM complex from the prospective and postmortem proteomic cohorts and from the RNA sequencing dataset on 443 melanoma tumor samples publicly available at the TCGA repositories. In each cohort, four groups were created based on the levels of the MCM2-7, where the lowest levels correspond to low proliferative tumors. The differential abundances of proteins and transcripts between higher and lower proliferative groups in each cohort were submitted to 2D functional annotation enrichment analysis using KEGG pathways and GO biological processes reported for the mitochondrial proteins: postmortem versus prospective cohorts **(B)**, prospective versus RNA sequencing cohorts **(C)**, and postmortem versus RNA sequencing cohorts **(D)**. Glutamate/glutamine **(E)**, reduced and oxidized glutathione ratios **(F)** and relative levels of S-formylglutathione **(G)** in the proliferation-based groups from the prospective cohort. **H)** Abundances of the catalase protein and transcript in the proliferation-based groups from the three patient cohorts. Significant differences are represented by *,**,***,****; q value < 0.05, 0.01, 0.001, 0.0001 respectively.

Proteins and transcripts from both the proteomic and transcriptomic cohorts were compared between the proliferation groups. The functional annotation enrichment analysis on dysregulated proteins and transcripts showed an overlap between the three cohorts. Pathways and processes linked to the metabolism of amino acids and lipids were over-represented by proteins and transcripts downregulated in the groups with higher tumor proliferation. The proteins of the mitochondrial translation machinery were jointly upregulated in highly proliferating melanomas from the different datasets (**Figure S7**). The relative levels of all mitochondrial transcripts and proteins between proliferation-based groups were submitted to a 2D functional annotation enrichment analysis (Cox and Mann, 2012). The resulting outcome could confirm that regardless of the sample origin, mitochondrial translation, RNA metabolic, and mitochondrial organization processes are upregulated in groups with higher proliferation (**Figure 3B-D**). On the other hand, biological pathways and processes linked to the metabolism of amino acids and lipids were commonly downregulated in the proliferating melanomas. The provided evidence highlights the dependence of melanoma tumor proliferation on mitochondrial function, which opens a therapeutic opportunity to target the mitochondrial translation machinery in the most aggressively proliferating melanomas. Moreover, these findings provided the foundations to deduce the mitochondrial function in melanomas by measuring the levels of transcripts or proteins of the MCM complex.

From the prospective cohort, the ratios between glutamate and glutamine were significantly larger in high proliferative group, suggesting that melanomas rely on glutamine consumption to promote their proliferation (**Figure 3E**). On the other hand, the GSH:GSSG ratios were lower in higher proliferative tumors, whilst higher levels of s-formylglutathione were observed (**Figure 3F-G**). This suggests that fast-growing melanomas are highly dependent on the glutathione metabolism. In addition to glutathione oxidation, catalase (CAT) also consumes hydrogen peroxide; however, the levels of catalase and its transcript in the three cohorts showed downregulation in the higher proliferative groups (**Figure 3H**). Altogether, these results indicate that melanomas, particularly those with a high proliferation rate, rely on glutathione rather than catalase to reduce the levels of ROS.

### 2.4. Mitochondrial proteome signature in melanoma metastases correlates with age and survival of patients

The relationships between the mitochondrial proteome and two clinical features of the patients; the age at diagnosis and the overall survival were explored. The proteins showing significant association with each clinical feature in the linear regression analysis were submitted to a functional enrichment analysis (**Figure 4A-B**). Among the proteins showing a positive association with the age of the patients, there were enriched biological pathways and processes linked to OXPHOS, TCA cycle, fatty acid degradation, the metabolism of amino acids as well as the urea cycle. Oppositely, signaling pathways and processes related to the immune system response and glycolysis showed negative correlations with age at diagnosis. Thus, melanomas from older patients seem to rely on mitochondrial pathways for their energy requirements, while proteins involved in triggering the release of cytochrome C from the mitochondria are downregulated, which ultimately drives apoptosis. The proteins showing significant correlation with the overall survival are involved in similar pathways to those associated with age; those showing a positive association with survival were negatively associated with age and vice versa. However, the mitochondrial translation machinery showed the most significant enrichment in proteins with a significant negative relationship with survival (**Figure 4B**). According to GLOBOCAN data, melanoma patients older than 60 have at least ten times higher mortality rate than younger patients (Ferlay et al., 2020). The current findings provide evidence on how the mitochondrial proteome of melanomas may be influenced by the age of the patient and how this can be linked to the aggressiveness of the tumor which ultimately results in shorter overall survival.

**Figure 4.**
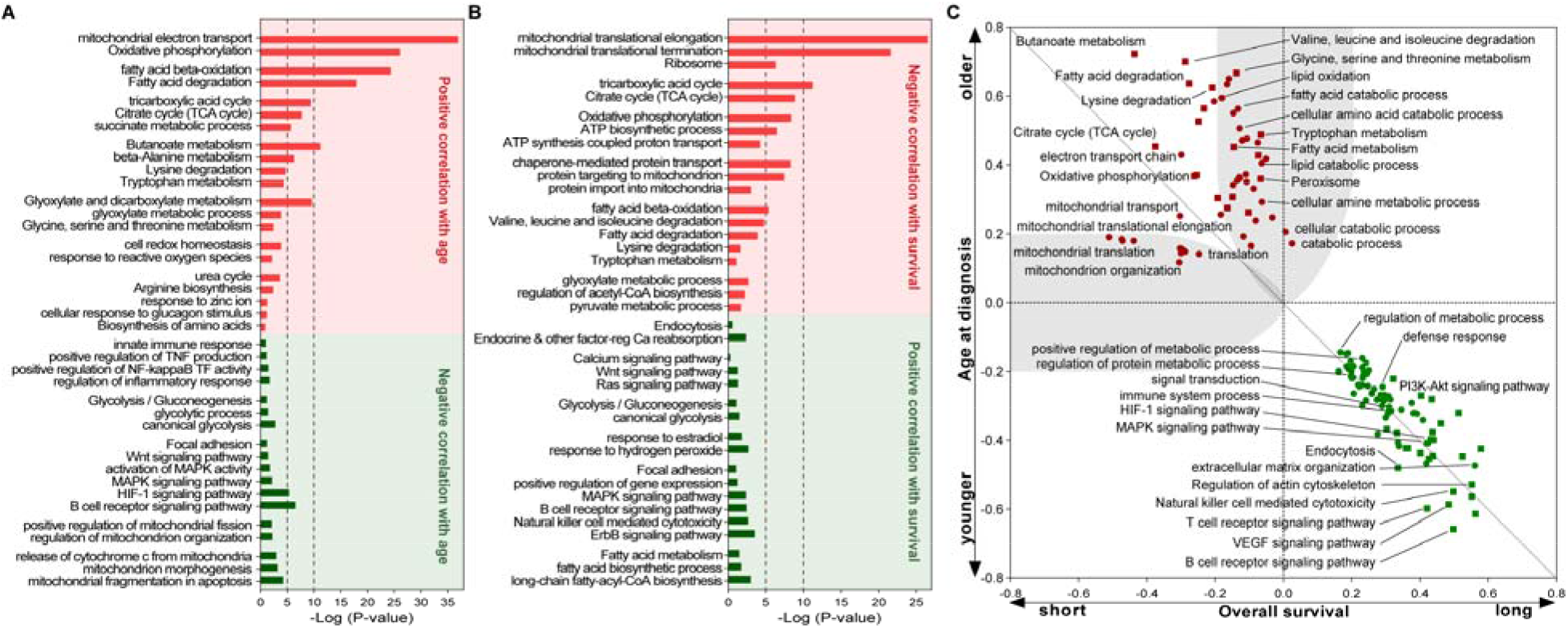
Functional enrichment analysis of mitochondrial proteins showing significant association with the age at diagnosis. **(A), and the overall survival of the patients (B)**. The analysis was performed using DAVID Bioinformatics. **C)** 2D functional annotation enrichment analysis using the slopes (beta) of the correlation between protein abundances and the overall survival of the patients (X-axis) and the age at diagnosis (Y-axis). The analysis was performed using the Perseus platform. Red and green dots correspond to biological annotations found significantly enriched in tumors from older and/or shorter survival patients, or younger and/or longer survival patients respectively.

The association between the mitochondrial proteome and the age at diagnosis and the overall survival was further investigated by 2D functional annotation enrichment analysis (**Figure 4C**). The analysis confirmed that similar pathways and processes were upregulated in melanomas from younger and long-term surviving patients, mostly linked to immune system response and to the regulation of metabolic processes. As previously pointed out, TCA cycle and OXPHOS showed a high positive association with both shorter survival and older patients. However, some biological annotations were more strongly associated with one of the clinical features. The mitochondrial translation machinery showed a stronger negative correlation with overall survival than its positive correlation with age. In contrast, most of the amino acid and lipid metabolism-related pathways showed a stronger association with age and have a lesser impact on the overall survival of melanoma patients.

### 2.5. Rewiring of mitochondrial pathways in metastases that arise after drug treatment

To gain insight into the contribution of mitochondrial pathways to the progression of melanoma following drug treatment, 44 melanoma metastases from the prospective cohort were analyzed. These tumors were divided into two groups, depending on whether the metastasis was developed and sampled before any therapy (naïve, n=26) or, whether it occurred during, or after the patient received treatment (treated, n=18). A multiple t-test analysis revealed 119 proteins significantly dysregulated between the two groups. The mitochondrial OXPHOS, mostly the complex I, the translation machinery, and to a lesser extent, some metabolic processes, were significantly enriched among proteins upregulated in metastases from the treated group (**Figure 5A**). On the contrary, processes related to the immune system response and regulation of apoptosis were enriched in naïve metastases. These findings suggest that there is a metabolic rewiring towards a more efficient way of maintaining the levels of energy production in metastases that developed after drug treatment.

**Figure 5.**
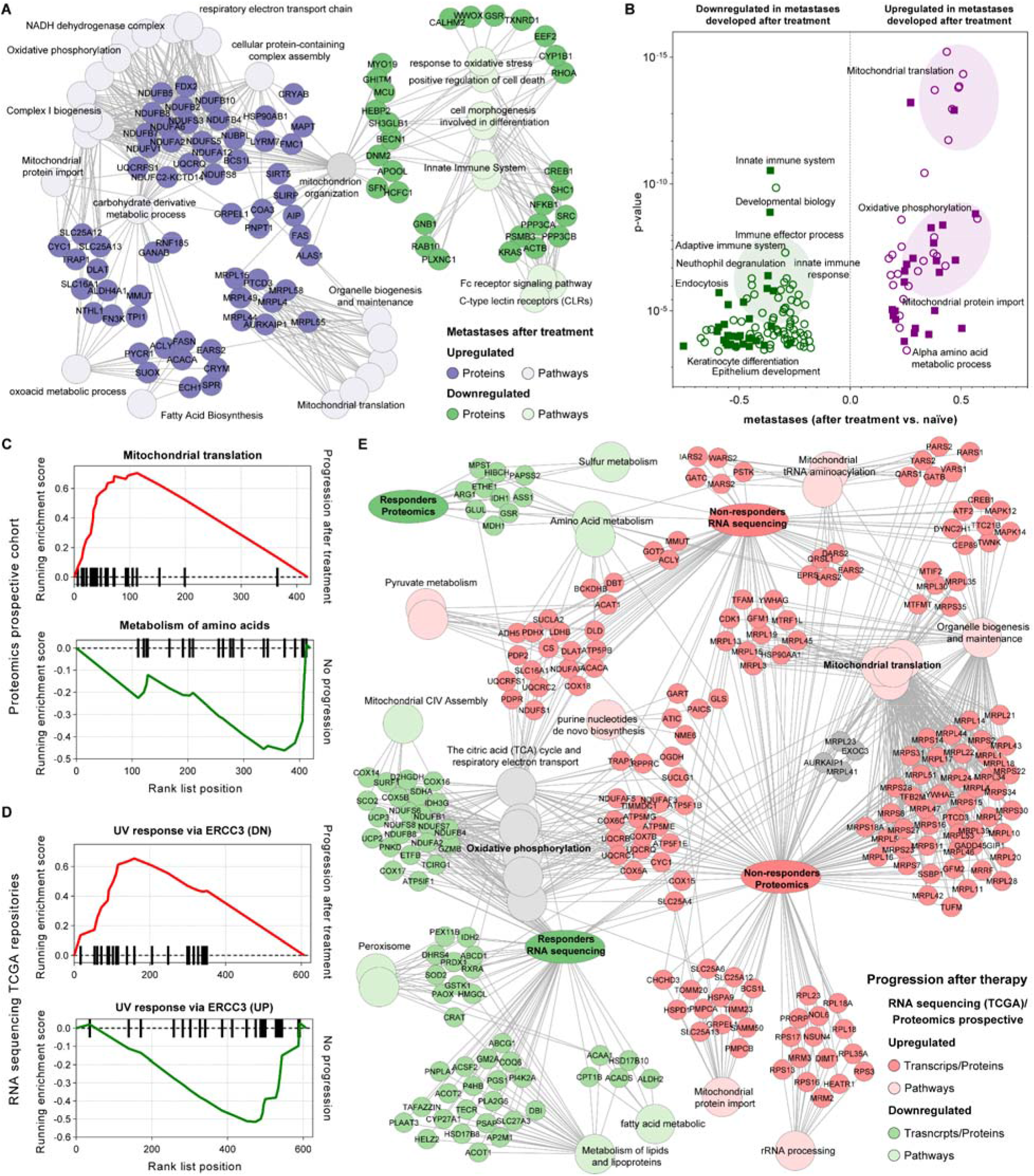
Mitochondrial translation machinery and OXPHOS upregulation have a strong impact on the progression of melanoma. **A)** ToppCluster enrichment analysis (Kaimal et al., 2010) and protein-pathway interaction network of mitochondrial proteins significantly dysregulated in metastases developed during or after the patients received treatment. **B)** Biological pathways (filled squares) and processes (circles) significantly enriched when comparing the mean protein abundances between the group of metastases that developed during or after treatment (purple) and those naïve (green). Data is represented as a volcano plot where the annotation enrichment p-value and the differences between groups were plotted. A false discovery rate (FDR) of 0.02 was set as the cut-off for significance. **C-D)** Gene Set Enrichment Analysis (GSEA) plots of pathways significantly enriched in tumors from patients where the disease progressed after they received treatment, **(C)** proteomics prospective cohort and **(D)** RNA sequencing dataset from the TCGA. **E)** ToppCluster enrichment analysis and protein-pathway interaction network of protein and transcripts significantly dysregulated between progression-based groups.

The relative abundances of the mitochondrial proteins between both groups of metastases were submitted to a 1D functional annotation enrichment analysis (Cox and Mann, 2012). The results confirmed that mitochondrial translation and the OXPHOS are the most significantly enriched biological annotations in the treated group. Furthermore, in this group of metastases proteins involved in the immune system response and signal transduction were significantly downregulated (**Figure 5B**). Together, these findings support the driver role of mitochondria during melanoma progression, producing energy and other intermediates while the immune responses are suppressed. Thus, mitochondrial OXPHOS and translation machinery may be explored as targets for treating metastases resistant to conventional therapy and suggest an opportunity to combine conventional treatment with mitochondrial inhibitors.

### 2.6. Mitochondrial translation upregulation suggests melanoma progression and poor prognosis after treatment

The mitochondrial proteome’s potential prognostic power to predict melanoma outcome was explored in both the prospective cohort and the RNA sequencing data from the TCGA. A total of 30 tumor samples from patients enrolled in the prospective study who relapsed or not after therapy were included. Following the same criteria, the transcriptome from 430 melanomas from the TCGA database, was also included. A gene set enrichment analysis (GSEA) revealed the mitochondrial translation as the most significantly enriched pathway in the group of non-responders, while the metabolism of amino acids was enriched in responders (**Figure 5C**). Similar analysis at the transcriptomic level revealed significantly enriched in non-responders, a group of genes downregulated in response to UV radiation including the mitochondrial translational initiation factor 2 (MTIF2). Oppositely, tumors from responders were enriched by genes upregulated because of UV radiation exposure including NDUFB7, NDUFS7, IDH2, and SCO2. These genes are involved in the TCA cycle and respiratory electron transport (Augusto Da Costa et al., 2005) (**Figure 5D**).

The differential abundance analysis in both cohorts revealed an output of 366 proteins and 527 transcripts significantly dysregulated between the groups of responders and non-responders to therapy. The joint functional enrichment analysis showed the mitochondrial translation machinery enriched in non-responders from both cohorts, highlighting its value for bad prognosis leading to disease progression. In addition, the TCA cycle and OXPHOS were enriched in the non-responders; however, at the transcript level a set of genes from the OXPHOS pathway were upregulated in tumors from responders to treatment (**Figure 5E**). For a good prognosis, at the transcript level genes involved in the metabolism of lipids and fatty acids, and at the protein level the metabolism of amino acids were enriched in the responder groups. Altogether, this joint analysis reinforces the hypothesis that high levels of the mitochondrial translation suggest an unfavorable prognosis and melanoma progression, as well as a novel direction for the development of potential therapeutic interventions.

### 2.7. BRAF mutation is associated with upregulation of the OXPHOS in melanoma

BRAF V600E is the most frequent driver mutation found in melanoma, accounting for about 50% of all primary tumors. Here, the possible influence of this mutation on mitochondrial function was explored. For this analysis, 57 samples from the prospective cohort were tested for BRAF mutation status; 22 tumors carried the wild-type (WT) variant while 24 and 11 carried the BRAF V600E and the BRAF V600K variants, respectively. From the TCGA RNA sequencing cohort, BRAF mutation status of 315 melanomas was previously reported; 151 carried the WT while 124 and 17 tumors carried the BRAF V600E and the BRAF V600K variants, respectively (Akbani et al., 2015). Quantitative proteomics on three melanoma cell lines, one carrying the BRAF V600E mutation (SK-MEL-28), and two carrying the BRAF wild type (SK-MEL-2 and VMM-1) were also included in the study. Differential protein or transcript abundance analyses between the groups were performed independently, followed by functional enrichment on the dysregulated gene products. The two BRAF WT cell lines were individually compared to the BRAF V600E cell line, and only those proteins commonly dysregulated relative to both WT cell lines were considered for further analysis. The results displayed a high degree of convergence in biological annotations, which were significantly enriched among the upregulated transcripts and proteins from BRAF V600E groups. The mitochondrial OXPHOS, together with pyruvate metabolism and TCA cycle were among the most enriched pathways and processes. Mitochondrial translation proteins were also found enriched in all BRAF V600E groups. However, downregulated proteins or transcripts in the BRAF V600E groups did not show any significant enrichment in biological pathways or processes.

Next, we investigated to what extent the most enriched biological annotations in all datasets represent a reliable shift of the set of proteins from each biological annotation. All proteins and transcripts reported to be involved in the pyruvate metabolism, TCA cycle, OXPHOS, and mitochondrial translation from the three datasets (prospective, RNA sequencing and cell lines) were submitted to the analysis. To compare the BRAF V600E group with the wild type, we performed paired t-tests using the differential abundance between the groups of the gene products involved in the same biological process (**Figure S8**). The OXPHOS pathway was found significantly shifted in all datasets toward the groups carrying the BRAF mutation. However, although elements of the mitochondrial translation were found upregulated in all datasets, when analyzed all together, in the RNA sequencing data, no significant shift to a particular group was observed. In contrast, at the protein level, all comparisons showed an upregulation of the mitochondrial translation in the BRAF V600E group relative to wild type (**Figure S8C**). A summary of the results from the proteomic analysis of tumors and cell lines, and transcriptomics from more than 300 melanomas, conclude that in BRAF V600E mutated tumors, the energy production is directed via metabolic changes towards the TCA cycle and the OXPHOS pathways. These findings further support the concept of a novel therapeutic strategy combining mitochondrial inhibitors with BRAF inhibitors.

### 2.8. Inhibition of mitochondrial ribosomes and OXPHOS affect melanoma cells

To further investigate the potential dependence of melanoma on mitochondrial function, we targeted mitochondrial pathways using established melanoma cell lines and non-cancerous melanocytes. The mitochondrial translation was targeted with three clinically approved antibiotics, two members of the tetracycline family (Doxycycline and Tigecycline) and the macrolide Azithromycin. Antibiotics from tetracycline and macrolide families, which reversibly inhibit the 30S and 50S subunits of bacterial ribosomes, also target their homologous subunits of the human mitochondrial ribosomes (Kim et al., 2017). Previously, these drugs exhibited anti-proliferative effects in several cancer stem cells (Hu et al., 2016; Lamb et al., 2015; Rok et al., 2020) and have been tested in clinical trials for treating different cancers (Chu et al., 2014; Ferreri et al., 2012; Reed et al., 2016). Three established melanoma cell lines (VMM-1, SK-MEL-2, and SK-MEL-28) in addition to HEMN-LP (non-cancerous melanocyte) were treated with the three antibiotics in doses ranging from 1.5 μM to 50 μM. The three drugs inhibited in a dose-dependent manner the proliferation of the melanoma cell lines; however, in all cases the non-cancerous melanocyte was more resistant to the treatment (**Figure 6A**).

**Figure 6.**
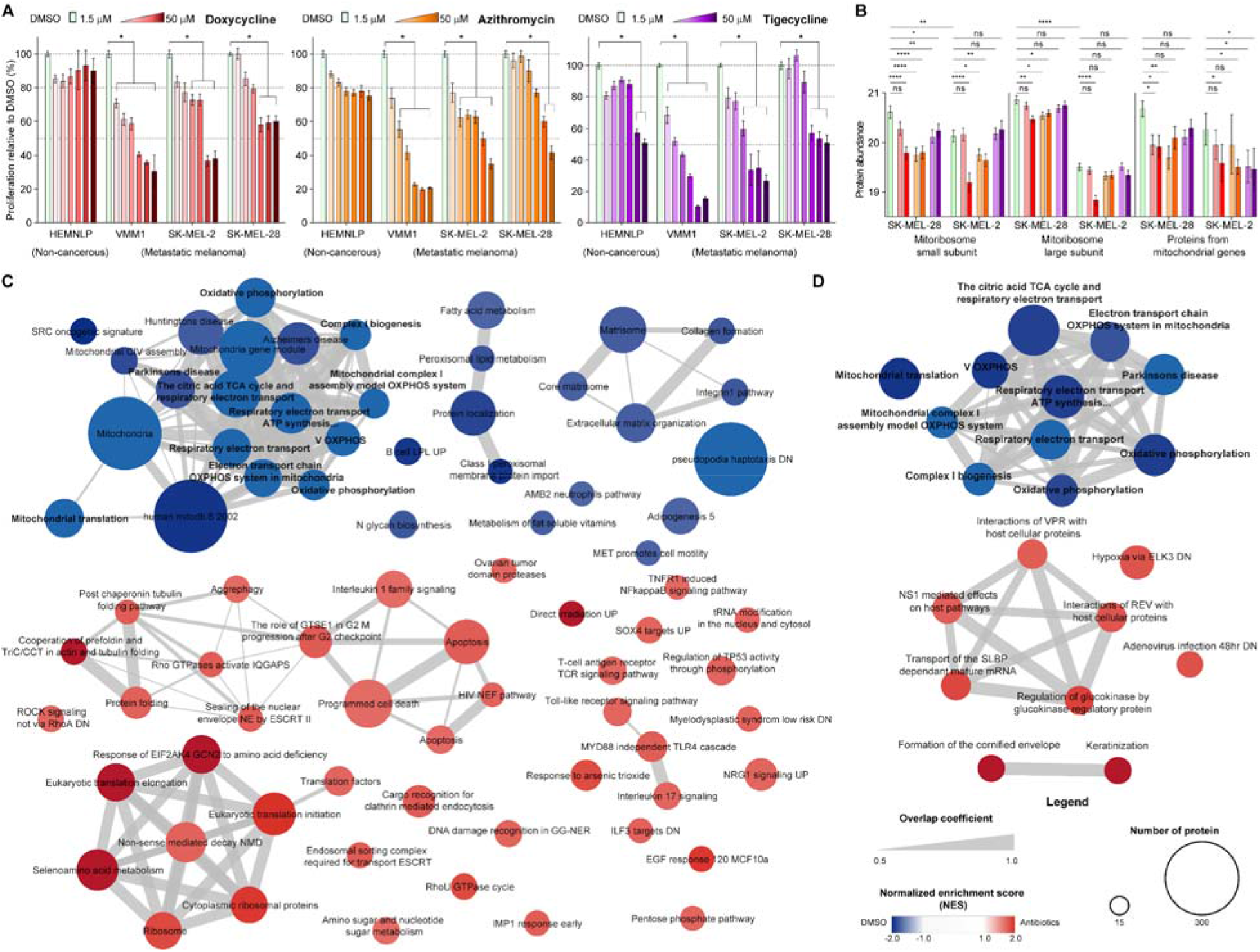
Anti-protein synthesis antibiotics target the mitochondrial translation and inhibit the proliferation of melanoma cell lines. **A)** Relative proliferation analysis in four different cell lines, HEMNLP (immortalized melanocytes, non-cancerous), VMM-1 (brain metastasis), SK-MEL-2 (cutaneous metastasis), and SK-MEL-28 (lymph node metastasis), treated with DMSO or antibiotics targeting the mitoribosomes. Doxycycline, Azithromycin, and Tigecycline were used at nine different doses in the low micromolar range (1.5, 3, 6, 12.5, 25, and 50 μM) and the proliferation was measured at 72 hours relative to DMSO treatment. Bars and errors correspond to the mean of three experimental replicates and the standard deviation respectively, * indicates significantly inhibited proliferation compared to DMSO treatment (p-value < 0.05, Kruskal-Wallis test and Dunn’s correction for multiple comparisons). **B)** Relative protein levels of the mitoribosome subunits and the protein products of the mitochondrial genes in SK-MEL-28 and SK-MEL-2 cell lines after 24 hours of treatment with two doses of the antibiotics. Data was extracted from the global proteomics analysis of the conditions tested. The statistical significance of the paired t-test between control-treated groups is represented (ns no significant, *, **, ***, and **** significantly different with p-value <0.05, <0.01, <0.001, and <0.0001 respectively). **C)** Pathways significantly dysregulated in SK-MEL-28 (**C**) and SK-MEL-2 (**D**) cell lines treated with antibiotics. The functional enrichment analysis was performed using the Gene Set Enrichment Analysis (GSEA) using all curated databases.

The mechanism-of-action of these antibiotics on the melanoma cell lines SK-MEL-28 and SK-MEL-2, each treated with vehicle or two different doses of the drugs for 24 hours, was explored through global quantitative proteomics. As expected, the inhibition of the mitochondrial translation machinery resulted in the downregulation of the proteins encoded by the mitochondrial genome and, to some extent, the mitochondrial ribosomes (**Figure 6B**). The proteome dynamics in response to the treatment with different antibiotics showed high similarity. The gene set enrichment analysis (GSEA) was performed considering all treatments for each cell line and summarized in **Figure 6C-D**. Commonly, antibiotics induce downregulation of the OXPHOS and the mitochondrial translation in both cell lines. SK-MEL-28 cells, which constitutively express higher levels of mitochondrial ribosomes, showed a more extensive set of pathways dysregulated. The treatment impaired the metabolism of fatty acids and pathways involved in the extracellular matrix organization and the integrin and collagen formation. This finding suggests that antibiotics, particularly in the more proliferative melanomas, not only inhibit the OXPHOS limiting the production of energy, but also affects the tumor microenvironment. On the contrary, the upregulated proteome was significantly enriched in apoptosis-related signaling pathways and the cytoplasmic ribosome. Altogether, we provided validity to the study by showing that antibiotics selectively impair the proliferation of melanoma cell lines through the inhibition of the mitoribosomes, which primarily results in the inhibition of the oxidative phosphorylation.

The metastatic melanoma dependence on the mitochondrial OXPHOS was explored by treating the above-mentioned cell lines (VMM-1, SK-MEL-2, SK-MEL-28, and HEMNLP) with three different compounds: VLX600, IACS-010759, and BAY 87-2243. All compounds were recently reported to target the mitochondrial OXPHOS (Ellinghaus et al., 2013; Molina et al., 2018; Zhang et al., 2014). These compounds have previously shown antitumor effect in human cancer models (Fryknäs et al., 2016; Gopal et al., 2019; Kanakkanthara et al., 2019; Karlsson et al., 2019; Mody et al., 2019; Schöckel et al., 2015; Vitiello et al., 2018). The three OXPHOS inhibitors (OXPHOSi) showed anti-proliferative effect on the examined melanoma cell lines in a dose-dependent manner (**Figure 7A**). VLX600 was anti-proliferative at the nanomolar to low micromolar dose range in melanoma cells while well tolerated by the non-cancerous cell line. The two other compounds were anti-proliferative at the micromolar range with higher sensitivity in melanoma cells compared to the non-cancerous melanocytes.

**Figure 7.**
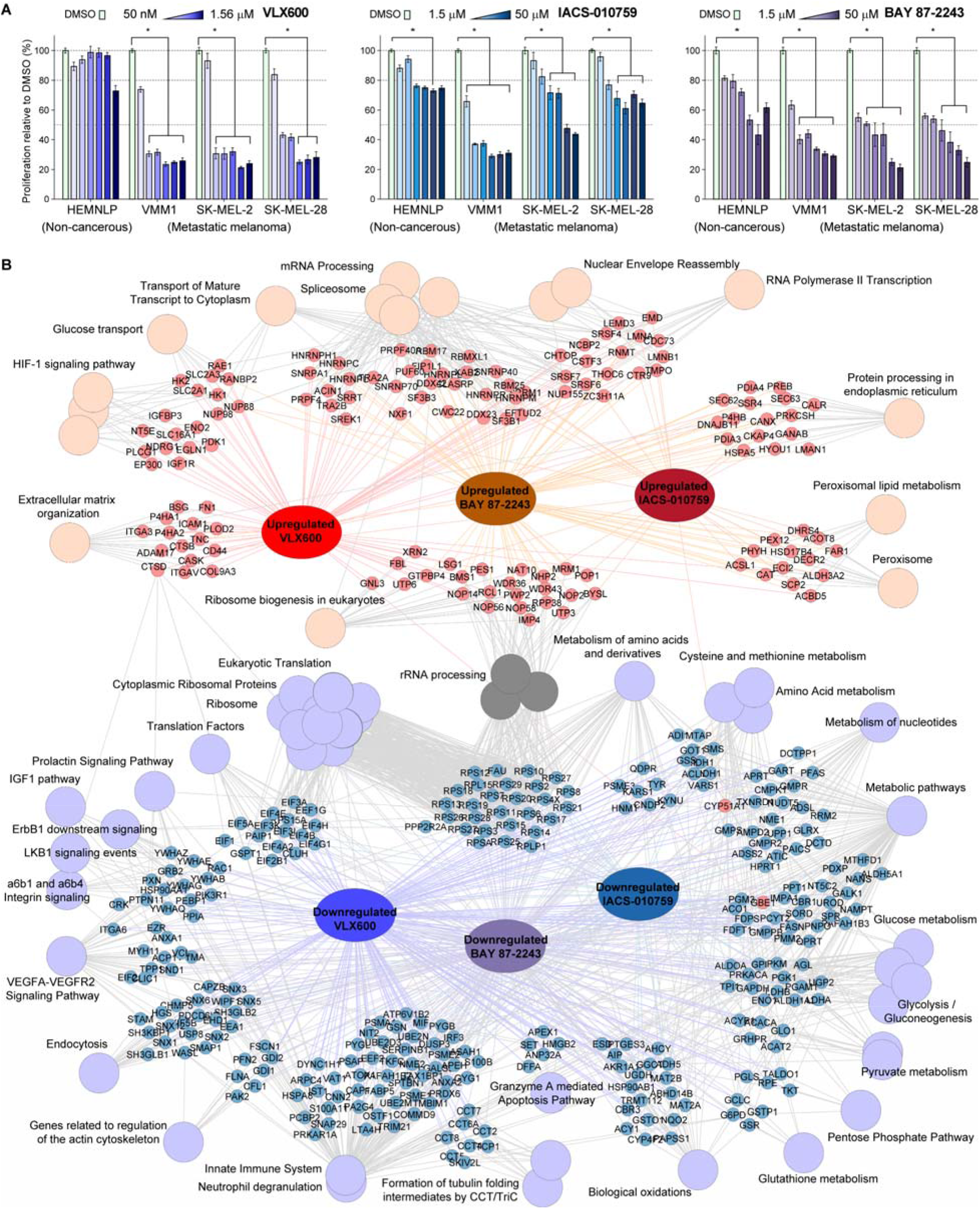
Drugs targeting the mitochondrial oxidative phosphorylation inhibit proliferation in melanoma cell lines. **A)** Relative proliferation analysis in four different cell lines, HEMNLP (immortalized melanocytes, non-cancerous), VMM-1 (brain metastasis), SK-MEL-2 (cutaneous metastasis), and SK-MEL-28 (lymph node metastasis), treated with DMSO or the OXPHOS inhibitors VLX600, IACS-010759, and BAY 87-2243. VLX600 was used at six different concentrations (50, 100, 200, 400, 800 nM, and 1.5 μM), IACS-010759 and BAY 87-2243 were evaluated in the low micromolar dose range (1.5, 3, 6, 12.5, 25, and 50 μM), the proliferation was measured at 72 hours relative to DMSO treatment. Bars and errors correspond to the mean of three experimental replicates and the standard deviation respectively, * indicates significantly inhibited proliferation compared to DMSO treatment (p value < 0.05, Kruskal-Wallis test and Dunn’s correction for multiple comparisons). **B)** Functional network interaction of the dysregulated proteome after treating the melanoma cell lines SK-MEL-28 and SK-MEL-2 with the OXPHOS inhibitors VLX600, IACS-010759, and BAY 87-2243. Each cell line was treated with two different doses of each drug and only proteins commonly dysregulated in both doses of the drug and in both cell lines were submitted to the functional enrichment analysis. The analysis was performed using the online version of ToppCluster function under the ToppGene suite. The provided curated pathways databases were used for the analysis and significant annotations were based on a p-value <0.05 with Bonferroni correction.

Global quantitative proteomics performed on two melanoma cell lines (SK-MEL-28 and SK-MEL-2) treated with vehicle or two different doses of the three OXPHOSi for 24 hours revealed key details of the mechanism-of-action of these drugs and the response of melanoma cells. The three drugs showed some common but also specific dysregulated proteins indicating slightly different mode-of-action. Commonly, these drugs downregulated the cytoplasmic translation machinery and its translation factors (**Figure 7B**). Metabolic pathways, including the glycolysis, pyruvate, and some amino acids metabolism were also commonly impaired by the three drugs. The glutathione metabolism and signaling cascades such as VEGFA-VEGFR2, ErbB1, and integrin signaling, frequently upregulated in cancers, were downregulated after treatment. Commonly, proteins involved in mRNA processing and splicing were found to be upregulated after treatment, similarly to the ribosome biogenesis and rRNA processing. Specifically, the treatment with VLX600 also triggered hypoxia-related signaling cascades (**Figure 7B**). Conversely, the cells respond to the treatment with BAY 87-2243 and IACS-010759 by upregulating the peroxisomal lipid metabolism and the protein processing in the endoplasmic reticulum. Altogether, these drugs impair the metabolism and the protein synthesis of melanoma cells, which ultimately result in the inhibition of proliferation.

Ultimately, we have provided in vitro data that support the melanoma dependence on mitochondrial translation and oxidative phosphorylation. By targeting these pathways with six different drugs, it was possible to selectively inhibit the proliferation of melanoma cell lines. In addition, global quantitative proteomics revealed important aspects of the mechanism-of-action of the drugs in the context of melanoma molecular pathology. These findings suggest a possibility for potential therapeutic intervention within the metastatic melanoma spectrum.

## 3. Discussion

Recently, several groups have pointed to the critical role that mitochondria play in certain types of cancers (Fulda et al., 2010; Porporato et al., 2018; Vasan et al., 2020; Weinberg and Chandel, 2015). Particularly, in melanoma, mitochondrial pathway dysregulations were linked to the ability to respond to immunotherapy (Harel et al., 2019). The commonly observed development of resistance to BRAF mutation targeted therapy has been associated with a metabolic switch from glycolytic to an OXPHOS phenotype (Trotta et al., 2017). The present study took a closer look at the dynamics of the mitochondrial proteome in a large cohort of melanomas, including corresponding tumor microenvironment and non-tumor samples. The major findings highlight the critical role of mitochondria in the development and progression of melanoma. The strong upregulation of mitochondrial translation and OXPHOS proteins observed in melanomas point to a functional dependence on these mitochondrial pathways. This is supported by high rate of assimilation of glutamine required for cancer cell proliferation (Choi and Coloff, 2019; Keshet et al., 2018; Nakagawa et al., 2009).

Mitoribosomes were earlier proposed and used as potential targets for anticancer therapies (Kim et al., 2017; Lamb et al., 2015; Škrtić et al., 2011). Recently, in an observational study on primary melanomas, mitochondrial translation upregulation was discovered in the disease recurrence group (Gil et al., 2021). In the present study, we observed positive correlations between the levels of mitoribosomes and the proliferation status of the tumors at both protein and transcript levels, as previously reported (Akbani et al., 2015). Also, higher levels of mitoribosomal proteins correlated with short survival and were found to be more abundant in metastases that developed during or after patients received treatment. The higher levels of mitoribosomes were indicative of a poor prognosis. Targeting the mitoribosomes with already approved antibiotics, intended to target bacterial protein synthesis, dose-dependently inhibited melanoma cells. An opportunity to develop targeted therapy for those tumors that do not respond to the first-line therapy is therefore suggested. The anticancer properties previously reported for these antibiotics in several cancer models can be concluded to be valid also for melanoma (Foroodi et al., 2009; Galván-Salazar et al., 2016; Lamb et al., 2015; Sun et al., 2009; Vendramin et al., 2021).

Melanomas in general, and specifically those from patients with shorter survival, showed an upregulation of the mitochondrial OXPHOS. The dependency on mitochondrial function was evaluated by inhibiting the OXPHOS in melanoma cell lines. Targeting the OXPHOS has previously been explored as an anticancer therapy in several cancer models including melanoma (Fulda et al., 2010; Hubackova et al., 2019; Jung et al., 2010; Martínez-Reyes et al., 2020; Trotta et al., 2017; Vasan et al., 2020; Weinberg and Chandel, 2015). The present data strongly indicate that melanomas rely, to a large extent, upon the mitochondrial OXPHOS. The mitochondrial ATP resultant from OXPHOS is exported to the cytoplasm via ADP/ATP transporters, also found upregulated. Particularly, ANT1 was reported to be highly expressed in oxidative cells (Hubackova et al., 2019). By using drugs inhibiting the OXPHOS, the metabolism and cytoplasmic translation of the cells are impaired, including signaling cascades that extend beyond tumor cells related to metastatic progression and angiogenesis.

In summary, the present study suggests a mitochondrial functional dependence in advanced melanoma. The upregulation of these critical mitochondrial pathways was accompanied by downregulation of immune response-related pathways. Likewise, observations made by Harel et al. highlighted that tumors from patients that responded to immunotherapy displayed upregulated mitochondrial pathways (Harel et al., 2019). There seems to be an evident change in the mitochondrial proteome during tumorigenesis and the progression of melanoma, which also extends beyond tumor cells into the adjacent microenvironment. Our results indicate that targeting the mitochondrial translation machinery and the OXPHOS may eliminate melanoma cells. Hence, a promising opportunity to treat metastatic melanomas, particularly useful for BRAF-mutated tumors, is presented. Although these tumors initially respond to BRAF inhibitors, many patients later develop resistance to the treatment and ultimately die. Targeting the mitochondria in melanomas, alone or in combination, may fill a gap in current melanoma therapy, especially for patients with a poor prognosis.

## 4. Experimental section

### Biopsy collections

#### Prospective cohort

Patients enrolled in the study turned with their primary cutaneous melanoma to the Department of Dermatology, Venereology and Dermatooncology, Semmelweis University, Budapest, Hungary. They underwent primary excision and native tumor tissue sampling was made by a dermatopathologist from the neighboring area of the thickest part of the tumor - not to hurt the correct diagnosis of the patient.

#### Postmortem cohort

22 individuals dead from melanoma underwent section and autopsy samples were collected from all the locations, where hematogenous metastasis could be detected. Together 74 solid organ samples from more than 20 different locations were investigated. Most of them were lung, liver, central nervous system, gastrointestinal and spleen sites.

### Ethical approval

The study was carried out in strict accordance with the Declarations of Helsinki and was approved by the Semmelweis University Regional and Institutional Committee of Science and Research Ethics (IRB, SE TUKEB 114/2012 and SE IKEB 191-4/2014). Samples were obtained from the Department of Dermatology and Venereology, Semmelweis University, Budapest, Hungary, under informed consent and a clinical protocol.

### Cell culture conditions and Reagents

All cell lines were validated and tested negative for mycoplasma (PCR Mycoplasma Test Kit, Promocell, Heidelberg, Germany). Cells were cultured in a humidified incubator at 37 °C and 5% CO_2_according to the recommendations and protocols of ATCC. SK-MEL-28 and SK-MEL-2 (ATCC, VA, USA) were cultured with Minimum Essential Medium (MEM, Corning, NY, USA) and 10% fetal bovine serum (Gibco, MA, USA) and 1% penicillin/streptomycin (HyClone, UT, USA). VMM1 (ATCC) were cultured with RPMI-1640 (Gibco) and 10% fetal bovine serum (Gibco) and 1% penicillin/streptomycin (Hyclone). HEMn-LP (Gibco) were cultured with Medium 254 (Gibco) supplemented with HMGS (Gibco). All cell lines were cultured according to the instructions supplied by the manufacturers.

### Cell proliferation assay

Following an established procedure (Wang et al., 2018), cell proliferation was assessed using a 3-(4,5-dimethylthiazol-2-yl)-2,5-diphenyltetrazolium bromide (MTT, VWR, PA, USA) assay. In brief, cells were seeded at 2 × 10^3^ cells/100 μl/well in 96-well plates and treated with or without drug. After incubation at 37°C for 24, 48, and 72 hr, MTT was added to each well, and cells were incubated at 37°C for 3 hr. The water-insoluble purple precipitate was then solubilized in 150 μl of Dimethyl sulfoxide (DMSO, Invitrogen) per well. The absorbance of the wells was measured at 595 nm with a reference wavelength of 650 nm using a Fluostar Omega Microplate Reader (BMG Labtech, Ortenberg, Germany).

Doxycycline hyclate (Sigma Aldrich), VLX600 (Sigma Aldrich), tigecycline (Sigma Aldrich), IACS-010759 (Cayman chemical), BAY 87-2243 (Sigma Aldrich), and Azithromycin dehydrate (Sigma Aldrich) were dissolved in DMSO (10 mM) and diluted in media for each concentration used. The concentrations used in the cell proliferation assay of doxycycline hyclate, tigecycline, BAY 87-2243, IACS-010759, and azithromycin dihydrate were 1.56 µM, 3.12 µM, 6.25 µM, 12.5 µM, 25 µM, 50 µM, 100 µM, and 200 µM. The concentrations of VLX600 were 48.83 nM, 97.66 nM, 195.31 nM, 390.63 nM, 781.25 nM, 1.5625 µM, 3.125 µM, 6.25 µM, and 12.5 µM, unless noted otherwise.

### Tissue processing

Frozen tissue samples were processed as previously described (Gil et al., 2019). Briefly, tissue pieces were sliced with a thickness of 10 μm to be used for histopathological characterization, proteomics, and metabolomics. For quantitative proteomics and metabolomics 15 to 30 consecutive slices were collected where the previous and immediately following slices were stained for computing the sample composition.

### Histological sample characterization

For histological characterization, frozen tissue sections (10 µm thickness) were placed on glass slides, stained with hematoxylin and eosin, and then scanned with an automated Aperio CS2 slide scanner system (Leica Biosystems, San Diego, CA, USA). The slides were then used to determine and evaluate the sample composition in terms of tumor cell, stroma, immune cell, adjacent tissue content. The annotations were performed utilizing the open-source software Qupath (Bankhead et al., 2017) (v0.2.0-m8) were selected/confirmed by a board-certified pathologist. After creating the region of interest (ROI) of each scanned tissue region, estimated stain vectors were collected by excluding unrecognized regions, such as white backgrounds in the Qupath software. In ROIs, tiles with separated squares of 5 µm x 5 µm were created, and from all square tiles, coherence features were analyzed, and detection classifiers were created by adding parameters from optical density, H&E, RGB OD, RGB, and Grayscale. The whole process was computed, each tumor sample was annotated by counting the separated squares, and number of pixels thereof, as their compositions were quantified at identical image sizes.

### Cell lines treatment for proteomics

Melanoma cells (SK-MEL-2 and SK-MEL-28) were seeded at 30 × 10^4^ cells/3 ml media/well in 6-well plates for 24 hr. Before treatment cells were in serum-starved conditions for 24 h. Cell were treated with DMSO or DMSO/drug for 24 h. The drugs were tested at two different doses each doxycycline hyclate (12.5 µM and 200 µM), Azithromycin (12.5 µM and 100 µM), Tigecycline (12.5 µM and 100 µM), VLX600 (390 nM and 6 µM), IACS-010759 (12.5 µM and 100 µM), and BAY 87-2243 (12.5 µM and 100 µM). After treatment, culture media were removed, and the cells were washed with PBS and stored at - 80 °C.

### Sample processing for proteomics

#### Prospective cohort and cell lines

A protein extraction buffer containing SDS 2%, DTT 50 mM, Tris 100 mM, pH 8.6, was added to the sliced tissues or cell pellet, rest for one minute and sonicated using a Bioruptor plus (40 cycles, 15 s On, 15 s Off, at 4 °C). Samples were incubated at 95 °C for 5 min. Proteins were alkylated by adding IAA to a final concentration of 100 mM for 20 min in the dark at room temperature (RT). The proteins were precipitated overnight by adding 9 volumes of cold ethanol. Proteins were dissolved in TEAB buffer containing SDS 1%, SDC 0.5%, and submitted to a chemical acetylation reaction of all free amino groups. The reaction was performed with N-acetoxy-succinimide-d3 (NAS-d3). O-acetylation was reverted by treating the samples with 5% hydroxylamine. Proteins were precipitated as described above to remove the excess reagents and dissolved in AmBiC containing SDC 0.5%. Proteins were digested with trypsin (1:50, enzyme:protein) at 37 °C, overnight. The SDC was removed from the peptide solution by adding ethyl-acetate and TFA (Gil et al., 2017a). After discarding the organic phase, the mixtures of peptides were quantified and kept at -80 °C until MS analysis.

#### Postmortem cohort

Tissue samples were mixed with a lysis buffer consisted of 100 mM ammonium bicarbonate and 4 M urea in an ice bath during 30 min. The samples were sonicated using a Bioruptor plus (40 cycles, 15 s On, 15 s Off, at 4 °C). The samples were centrifuged at 10,000g for 10 min at 4 °C, and the protein content was determined using the colorimetric micro–BCA Protein Assay Kit according to the instructions supplied by the manufacturer (ThermoFisher Scientific, Rockford, IL). The AssayMAP Bravo platform (Agilent Technologies) was used to perform urea in-solution digestion as published before (Kuras et al., 2019). Briefly, 40-50 µg of proteins were processed for each sample using the In-Solution Digestion protocol: Single Plate v1.0 protocol in VWorks. The samples were first reduced with 10 mM DTT for 1 h at 37 °C and alkylated with 20 mM iodoacetamide for 30 min in the dark at RT. This was followed by enzymatic digestion with endoproteinase Lys-C in a 1:50 w/w ratio (enzyme/protein) in 1 M urea and incubated at RT for 7 h. Next, the samples were diluted with 100 mM ammonium bicarbonate to a concentration of 0.6 M urea, trypsin was added in a 1:50 w/w ratio (enzyme/protein) and the digestion was incubated overnight at RT. The peptides generated were acidified by adding 4 μL of 50% formic acid and desalted using the Peptide Clean-up v2.0 protocol. Here, C18 cartridges (Agilent, 5 μL bead volume, 150 μg capacity) were primed using 100 μL of 90% acetonitrile and equilibrated with 70 μL of 0.1% TFA at 10 μL/min. Samples were loaded at 5 μL/min, followed by internal cartridge wash and cup wash with 0.1% TFA at 10 μL/min. The peptides were eluted using 30 μL 80% ACN with 0.1% TFA at 5 μL/min and stored dry at -80 °C until MS analysis.

#### Cell lines with treatment

A protein extraction buffer containing SDS 5%, DTT 20 mM, TEAB 50 mM, was added to the cell pellet, rest for one minute and sonicated using a Bioruptor plus (30 cycles, 15 s On, 15 s Off, at 4 °C). Samples were incubated at 95 °C for 5 min. Proteins were alkylated by adding IAA to a final concentration of 50 mM for 20 min in the dark at room temperature (RT). Samples were loaded on a S-Trap 96 well plate (Protifi) for protein digestion following manufacturers’ instructions. Proteins were digested with trypsin (1:50, enzyme:protein) at 37 °C, overnight. Peptides were eluted from the S-Trap plate by adding three consecutive volumes of digestion buffer (50 mM AmBiC), 0.2% aqueous formic acid, and 0.2% formic acid in ACN/H2O 1:1 mixture, follow centrifugation after each addition. Peptides were dried and kept -80 °C until MS analysis.

### Protein identification and quantification

#### Peptide spectral library

Pools of peptides including all the samples from the prospective, postmortem, and cell line cohorts enrolled in the study were spiked with iRT and analyzed in a high-resolution LC-MS system (Dionex Ultimate 3000 RSLCnano UPLC coupled to a Q-Exactive HF-X mass spectrometer (Thermo Fischer Scientific)). Peptides were desalted on a trap column Acclaim PepMap100 C18 (3 µm, 100 Å, 75 µm i.d. × 2 cm, nanoViper) and then connected in line with an analytical column EASY-spray RSLC C18 (2 µm, 100 Å, 75 µm i.d. × 50 cm). The temperature of the trap column and analytical column were set at 35 °C and 60 °C respectively, and the flow rate was 300 nL/min. For the library building of the prospective cohort, peptides were separated using a non-linear gradient of 137 min, from 2 to 27% of solvent B in 112 min, followed by increasing from 27 to 35% in 10 min, then to 55 % in 5 min, then to 90% in 5 min, and finally maintained at 90% of B for 5 min. For the postmortem cohort the gradient was slightly different, the duration was 137 min and went from 2 to 25% of solvent B in 112 min, followed by increasing from 25 to 32% in 10 min, then to 45 % in 5 min, then to 90% in 5 min, and finally maintained at 90% of B for 5 min. Data were acquired applying a top 20 data-dependent acquisition (DDA) method over a mass range of 385-1460 m/z. The parameters for MS1 included a resolution of 120,000 (@ 200 m/z), the target AGC value of 3E+06, and the maximum injection time was set at 100 ms. For MS2, the resolution was 15,000, the AGC value 1E+05, the injection time 50 ms, the threshold for ion selection 8E+03, and the normalized collision energy (NCE) 28. Selected ions were excluded for 40 s and single charged ions were not eligible for fragmentation. The isolation window was 1.2 Th. The cell lines submitted to treatment were analyzed in the same system and acquired using a DDA method as mentioned above. The peptides were separated in a non-linear gradient of 63 minutes, from 4 to 7% of B in 2 min, from 7 to 28% in 50 min, followed by increasing to 40% in 5 min, then to 55% in 5 min, and finally to 90% in 1 min.

#### DIA analysis

individual samples spiked with iRT were analyzed in duplicates using the same LC-MS system and with the same chromatographic conditions as for building their library. Data were acquired following a Data Independent Acquisition (DIA) method. The method was intended for peptide and protein quantification using the MS1 scans. The design of the DIA method is composed of a full MS cycle containing 54 variable width MS2 windows and 3 full scan MS1. A full MS1 scan was placed every 18 MS2 scans. The width of the MS2 windows were selected based on the empirical m/z distribution of identified peptides for the spectral library building. Between adjacent MS2 windows, an overlap 1 Da was placed. The variable windows were slightly different in each cohort depending on the identified peptides in the library. The parameters for MS1 were resolution 120,000 (@200 m/z), AGC target 3E+06, injection time 50 ms, range 385-1460 Th. For the MS2 were resolution 30,000, AGC target 1E+06, injection time 50 ms, NCE 28.

Protein and peptide identification and quantification were performed with the Spectronaut software (Biognosys, AG). DDA raw files coming from samples and pools processed with the same workflow were searched together, against a human protein database downloaded from UniProt in 2018, to create a spectral library. The search engine used was pulsar, which is integrated into the Spectronaut platform. For the prospective cohort, the parameters included Arg-C as the cleavage enzyme, 2 missed cleavages were allowed, lysine acetylation (d3) and carbamidomethyl-cysteine were set as fix modifications, methionine oxidation and protein N-terminal acetylation (d3) and (d0) were set as variable modifications. For the postmortem cohort, the selected enzyme was trypsin, two missed cleavages were allowed, carbamidomethyl-cysteine was fixed, and methionine oxidation and protein N-terminal acetylation were set as variable. The identifications were controlled by an FDR of 1% at the PSM, peptide and protein levels. The generated library for the prospective cohort included 200,000 different peptides from 13,000 proteins, while the library used in the postmortem cohort contains 300,000 peptides and 13,500 proteins. The DIA runs were calibrated using the iRT, and the identifications were defined by q-value cutoff of 0.01. The peptide and protein quantification were based on the intensity of the precursors.

### Amino acids and metabolites profiling

Twenty-one amino acids and some metabolites were extracted from frozen sliced tissue (∼ 7.8 mg) with ice-cold methanol spiked with 50 nM Testosterone-D3 (National Measurement Institute, Australia). The samples were sonicated for 30 minutes at 30 °C ± 2, then centrifuged at 14,000 g for 15 min at 4 °C. The supernatants were collected, dried (SpeedVac ™, Thermo Scientific), and reconstituted in 0.1% formic acid (Bobeldijk et al., 2008). A pool of all samples was used as quality control and blank samples (0.1% formic acid) were used eliminate potential contamination. For the amino acid profiling, samples were diluted 10 times with a combination solvent of 1:2:1 (v/v/v) H2O/ACN/MeOH supplemented with stable isotopic labeled amino acids solution at 1 nmol/mL (MSK-A2-1.2, Cambridge Isotope Laboratories, USA).

#### Amino acid profiling

Each sample was analyzed in triplicate by direct flow injection analysis (FIA) (Dionex Ultimate 3000 UHPLC, Thermo Scientific) connected to a Q-Exactive Plus (Thermo Scientific). The mobile phase adopted was H2O/ACN/MeOH 1:2:1 (v/v/v) and flow rate set for 50 μL/min. All amino acids were detected in positive mode by electrospray ionization and Target-SIM data acquisition. Mass spectrometry parameters were set as follows: capillary temperature, 280 °C; gas temperature, 150 °C; spray voltage, 3.50 kV; 30 sheet gas; 10 auxiliary gas; resolution 70 000; AGC target 5e5; isolation window, 1.0 m/z; acquisition time, 2 min. All parameters associated with sample dilution, FIA, ionization, and mass analyzer were optimized to obtain a symmetric peak with precision and reproducibility. Data were analyzed by Tracefinder Forensic software (Thermo Scientific). The values reported for each sample represents the ratio of peak intensity for the amino acid vs. its internal standard.

#### Metabolite profiling

8 µL of each sample was injected in triplicate and analyzed in the positive and negative mode in a LC-ESI-MS system (Dionex Ultimate 3000 UHPLC and Q-Exactive Plus, Thermo Scientific). Chromatographic separation was performed using a reverse phase column (C18 Zorbax, 50 × 3 mm, 1.7 μm, Agilent) and a mobile gradient phase (400 μL/min) composed of solution A (0.1% formic acid and 5mM ammonium formate) and B (methanol acidified with 0.1% formic acid). Gradient mobile phase was employed over 20 min, which was distributed as follows: 0-1 min, 5% B; 1-14 min, linear gradient from 5 to 100% B; 14-16 min, 100% B; 16-16.5 min, decreasing linear gradient from 100% to 5% B; 16.5-20 min, 5% B. The column was maintained at 40 °C, and the sample chamber was held at 7 °C. The samples were analyzed in full MS mode (m/z 70-800) with data-dependent acquisition (dd-MS2, top-10 DDA), normalized collision energy of 30, and MS1 and MS2 resolution of 70 000 and 17 500, respectively. Source ionization parameters were set as follows: capillary temperature, 380 °C; auxiliary gas temperature, 380 °C; spray voltage, 3.90 kV (positive mode) and 2.90 kV (negative mode); 60 sheath gas and 20 auxiliary gas. Data were analyzed by software Compound Discoverer 3.1. Metabolite identifications were based on parent and fragment ions and supported by comparison with public databases (Human Metabolome Database (HMDB), Kyoto Encyclopedia of Genes and Genomes (KEGG), Lipid Maps®, BioCyc Database Collection, and MZcloud). It was accepted a mass measurement accuracy of 5 ppm (Monnerat et al., 2019).

### Mitochondrial proteins filtering and normalization

The protein quantification output in each dataset was processed under Perseus platform (Tyanova and Cox, 2018). Values were log2 transformed and subtracted by the median in each run. The abundance of the proteins in the two replicates were averaged. The mitochondrial proteins were filtered based on the report for mitochondrial location in MitoMiner v4.0 (Smith and Robinson, 2019). The total list of mitochondrial proteins reported in the repositories of MitoMiner was 2504 of which, 1713 and 1911 were identified in the prospective and postmortem cohorts, respectively. In the case of the prospective cohort 1287 proteins and the postmortem cohort 1537 proteins were quantified in at least 70% of the samples, which represents the 75% and 80% of all identified proteins, respectively. The identified proteins from cell lines included 1620 proteins classified as mitochondrial. Filtered mitochondrial proteins were then subtracted by the median in each sample to correct for potential differences in the level of mitochondria. No imputation of missing values was imposed to any of the datasets under study.

The dynamics of the mitochondrial proteome in the prospective cohort relative to the origin of the samples was explored through an ANOVA (FDR 0.05) and subsequence post-hoc, under Perseus platform. The multiple t-test for two group comparisons between sample origin groups were performed in GraphPad. For the analysis, the variance was assumed at the level of individual protein, and the FDR, based on the method Two-stage step-up (Bejamini, Kriege, and Yekutieli) (Benjamini et al., 2006), was set at 5%.

The levels of proteins and transcripts of the MCM complex elements were extracted after total proteins and transcripts normalization and averaging the sample replicates. Hierarchical clustering analyses of the samples were performed using the relative levels of the six elements of the complex (MCM2-7). The cluster distances were computed using the Euclidian distance, and the method was ward.D. The analyses were done in R. In the three cohorts four groups were created based on the MCM complex levels, where the groups of samples with lower levels of the proteins or transcripts were tagged as Low proliferative, and the groups with the highest levels were tagged as High proliferative. Middle groups were labeled as Medium-Low and Medium-High following the same logic. The differential expression analyses between proliferation groups in the prospective, postmortem and RNA sequencing cohorts were performed through multiple t-test. The FDR of 5% was set, based on the method Two-stage step-up (Bejamini, Kriege, and Yekutieli). The association between the mitochondrial proteome and the proliferation of melanomas was assessed through 2D functional annotation enrichment (Cox and Mann, 2012) analyses between the different cohorts. The differences between higher and lower proliferative groups were averaged in each cohort, and these values were used for the enrichment analysis using KEGG pathways (Kanehisa et al., 2017) and GO biological processes (Carbon et al., 2017) between the cohorts.

The relationships between mitochondrial proteome and the age and overall survival of the patients, in the postmortem cohort, were analyzed by doing a linear regression test per protein in R. The p-values were adjusted to control the FDR provoked for multiple testing and a FDR up to 5% was accepted. Proteins with significant positive or negative association with the age or overall survival of the patients were used for functional enrichment analysis on biological pathways and processes using the online version of DAVID bioinformatics (https://david.ncifcrf.gov/). The slopes for each protein and both clinical feature associations were loaded in Perseus and submitted to a 2D functional annotation enrichment analysis using default settings (Cox and Mann, 2012). The biological annotation for each protein was taken from KEGG pathways and GO biological processes.

Metastases in the prospective cohort were grouped based on their treatment status: one group included 26 metastasis tumors that were taken before the patients received any drug treatment (naïve), the other group was composed by metastasis tumors taken during or after patient received treatment (treated). A multiple t-test between both groups was performed in GraphPad. The variance was assumed equal at the level of individual protein, and the cutoff p-value for significant differences was set at 0.05, without correction for multiple comparisons. Additionally, the individual protein difference between the average in each metastasis group (naïve and treated), were taken as the input for performing a 1D functional annotation enrichment analysis under Perseus platform (Cox and Mann, 2012). The functional protein annotation was taken from the curated dataset reported by GSEA (Mootha et al., 2003; Subramanian et al., 2005) and GO biological processes. Significantly enriched annotations were controlled by applying an FDR of 0.02 in Perseus software.

In the prospective cohort, for 30 of the tumor samples the information regarding the progression of the disease after therapy was collected. A total of 9 tumors came from patients which positively responded to therapy and 21 tumors were from patients that relapsed after receiving therapy. In addition, the RNA sequencing cohort was included in the analysis. For 430 tumors, the same information regarding the progression of the disease after treatment was available, 190 samples came from patients in which the disease did not progress, and 240 samples were from patients that relapsed after therapy. Both datasets were submitted to a multiple t-test for comparing between progression-based groups, performed in GraphPad. The variance was assumed equal at the level of individual protein or transcript, and the cutoff p-value for significant differences was set at 0.05, without correction for multiple comparisons.

The association of the BRAF mutation status, particularly BRAF V600E, of the tumors and mitochondrial proteome was investigated in the prospective cohort, the RNA sequencing cohort and in three melanoma cell lines. In the prospective cohort, the BRAF mutation status was assessed for 57 tumor samples, 22 tumors contain the WT variant of BRAF, 24 carry the BRAF V600E mutation, and 11 carry the BRAF V600K mutation variant. For the RNA sequencing cohort, the BRAF mutation status was reported for 315 tumors (Akbani et al., 2015). A total of 151 tumors were BRAF WT, 124 carried the BRAF V600E mutation, 17 carried the BRAF V600K, and 23 carried other less common BRAF mutations. The influence of the BRAF V600E mutation on the mitochondrial proteome was investigated through multiple t-test analyses between the groups carrying the mutation and those with the WT variant of the BRAF gene. The protein and transcript elements of several mitochondrial pathways, such as pyruvate metabolism and TCA cycle, the OXPHOS, and the mitochondrial translation, were submitted to paired t-test to compare the means of the BRAF WT and BRAF V600E groups in each cohort. The average of the differences between each of the pairs was considered significantly shifted from zero when the two-tailed p-value was < 0.05.

The response of the metastatic melanoma cell lines to the antibiotics and OXPHOSi was explored through global proteomics and subsequent bioinformatics analysis. A total of 6561 proteins were identified, including 6016 quantified in 70% of the samples. Protein abundances were normalized as mentioned above. Multiple t-test analyses between each treatment and their corresponding control were performed. The cut-off for considering significantly dysregulated proteins was set to accept a FDR up to 5%. Significantly dysregulated proteins were submitted to a functional enrichment analysis using the online version of ToppCluster (https://toppcluster.cchmc.org/) (Kaimal et al., 2010). Cells treated with antibiotics were analyzed using the GSEA software (Subramanian et al., 2005). The visualization of the results was performed in Cytoscape v3.9 (Merico et al., 2010).

## 5. Acknowledgments

This study was supported by the Berta Kamprad Foundation, Thermo Fischer Scientific and by grants from the National Research Foundation of Korea, funded by the Korean government (2015K1A1A2028365 and 2016K2A9A1A03904900), the Brain Korea 21 Plus Project, Republic of Korea, and ICONS (Institute of Convergence Science), Yonsei University, Republic of Korea. This work was done under the auspices of a Memorandum of Understanding between the European Cancer Moonshot Center in Lund and the U.S. National Cancer Institute’s International Cancer Proteogenome Consortium (ICPC).

## Author Contributions

J.G., Y.K., M.K. and L.H.B. designed and performed experiments. J.G. and Y.K. analyzed the data. J.G. draft the manuscript. J.G., E.W., J.M., K.P. and G.M.-V. wrote the paper. V.D., U.C., R.A., H.O., B.L., J.T.,S.K., and G.M.-V. collect the patient samples. J.G., Y.K., LH.B., A.S., R.A., F.CS.N., G.B.D., J.M., B.B., E.W., I.B.N., A.M.S., H.J.K., K.P., D.F., J.T., M.R., S.K., and G.M.-V. provided the scientific oversight of the study. Y.K., Y.S., H.O., B.L., I.B.N., and A.M.S. generated and analyzed histological images. J.G., I.P.P., R.H., and D.F. performed bioinformatic analyses. J.S.G., G.M., G.R.A.C., F.C.S.N. and G.B.D., designed and performed the metabolomics and amino acid analysis.

## 7. Supplemental figures

**Figure S1.**
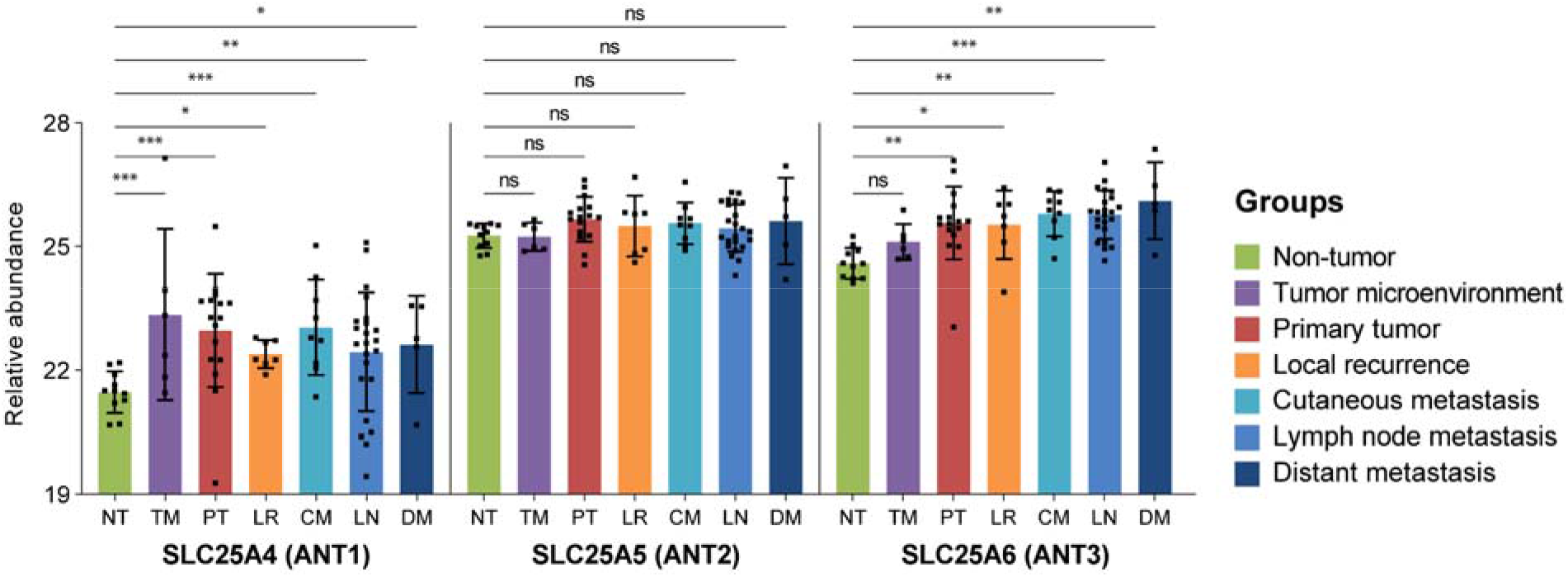
ADP/ATP translocases 1 and 3 are upregulated in melanomas and their microenvironment. Bars correspond to the median abundance in the group and the error lines indicate the standard deviation. *, **, *** indicate significant differences compared to the non-tumor group (q-value < 0.05, 0.01 and 0.001 respectively), ns indicates not significant.

**Figure S2.**
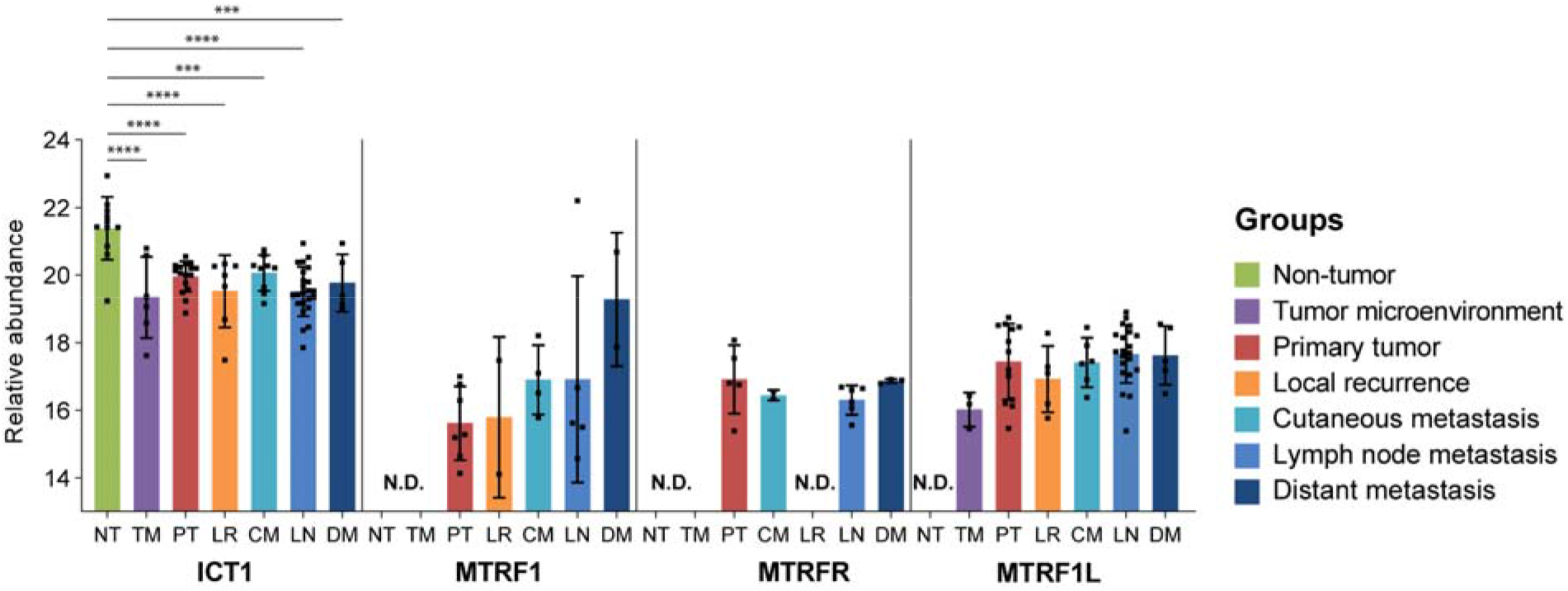
Relative abundance profiles of the four members of class 1 releasing factors of the mitochondrial protein synthesis. ***, **** indicate significant differences compared to the non-tumor group (q-value < 0.001, 0.0001 respectively)

**Figure S3.**
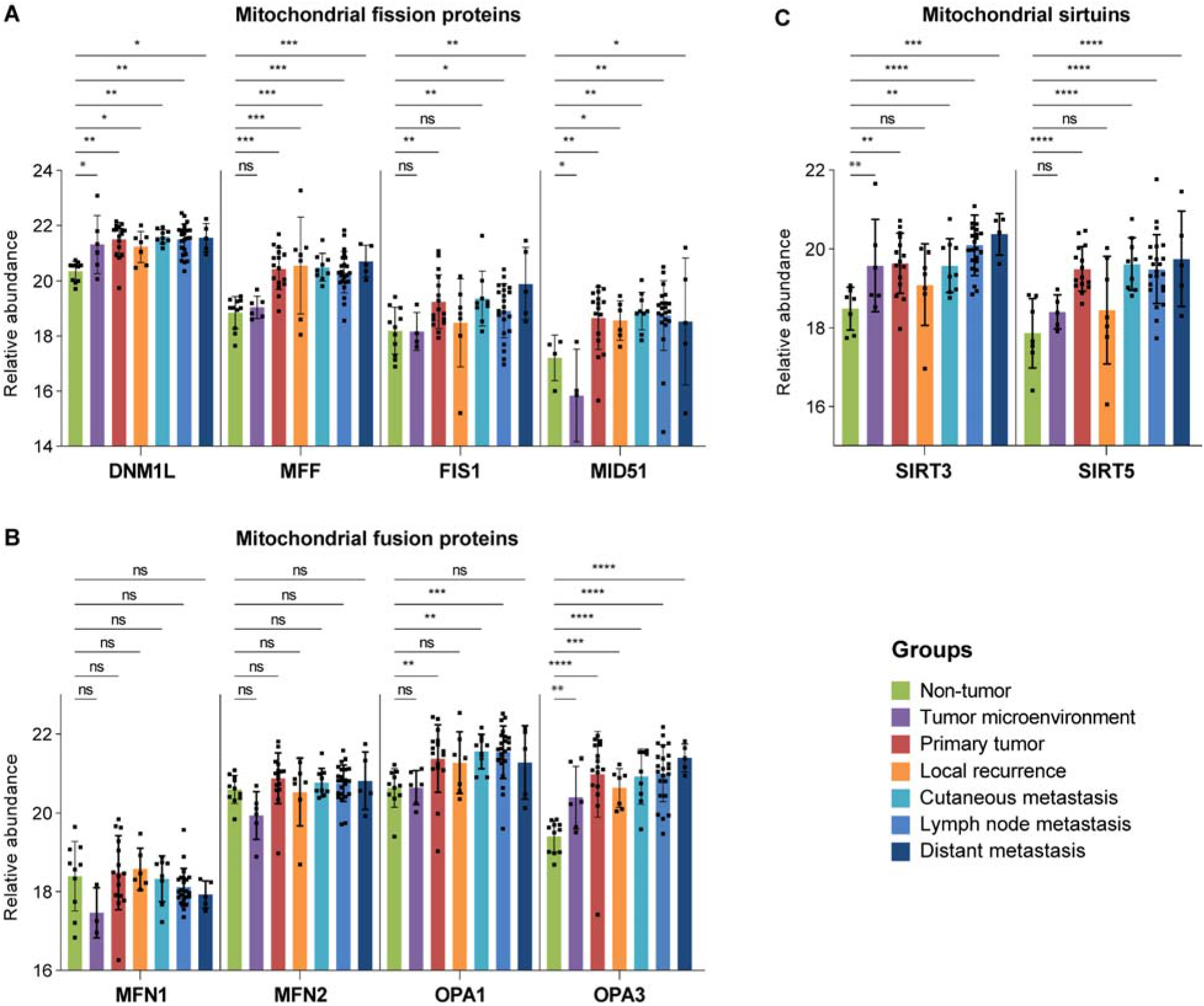
Mitochondrial dynamics in melanoma based on the sample origin. Relative abundance profiles of proteins involved in mitochondrial fission **(A)** and mitochondrial fusion **(B)** processes, grouped based on sample origin. **C)** Mitochondrial sirtuins (SIRT3 & SIRT5) relative abundance levels in melanoma-related groups of samples. Bars indicate the mean intensity of the protein in each group, error bars represent the standard deviation and the color of the group based on the origin of the sample (see legend). *, **, ***, **** indicate significant differences compared to the non-tumor group (q-value < 0.05, 0.01, 0.001 and 0.0001 respectively), ns indicates not significant.

**Figure S4.**
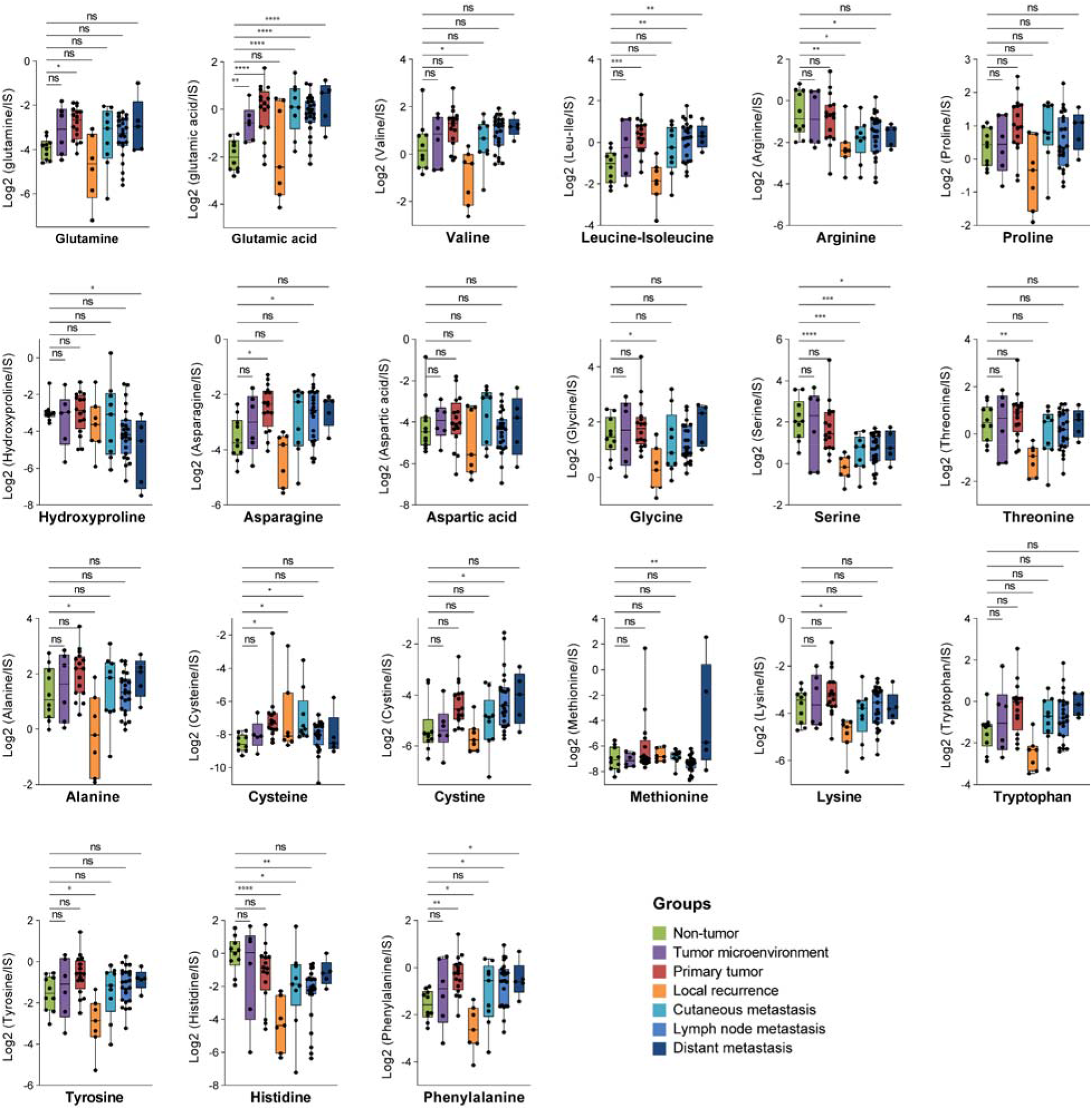
Abundance profile of free amino acids in melanoma-related tissues from the prospective cohort. Quantitative values are reported grouped based on the sample origin, Non-tumor, Tumor microenvironment, Primary tumor, Local recurrence, Cutaneous, lymph node, and distant metastases. The comparison between non-tumor and all other groups was performed with One-way ANOVA following an FDR approach. *, **, ***, **** indicate significant differences compared to the non-tumor group (q-value < 0.05, 0.01, 0.001 and 0.0001 respectively), ns indicates not significant.

**Figure S5.**
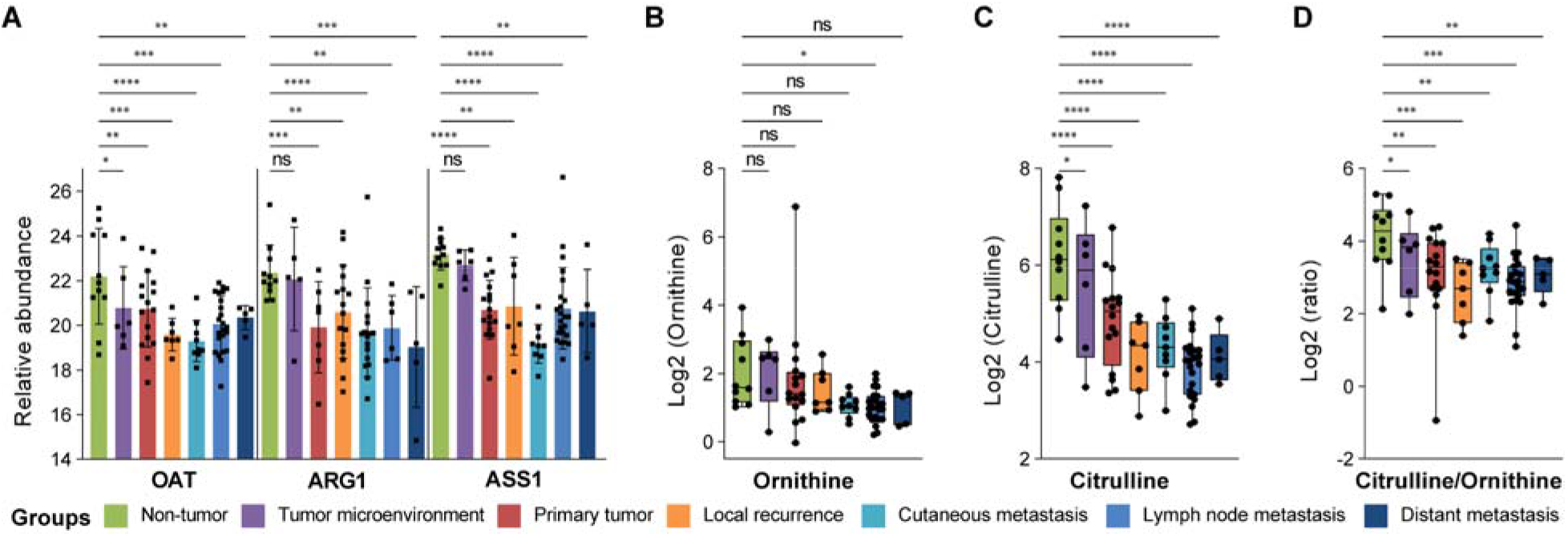
Relative levels between sample origin groups in the prospective cohort of enzymes and metabolites involved in the urea cycle. *, **, ***, **** indicate significant differences compared to the non-tumor group (q-value < 0.05, 0.01, 0.001 and 0.0001 respectively), ns indicates not significant.

**Figure S6.**
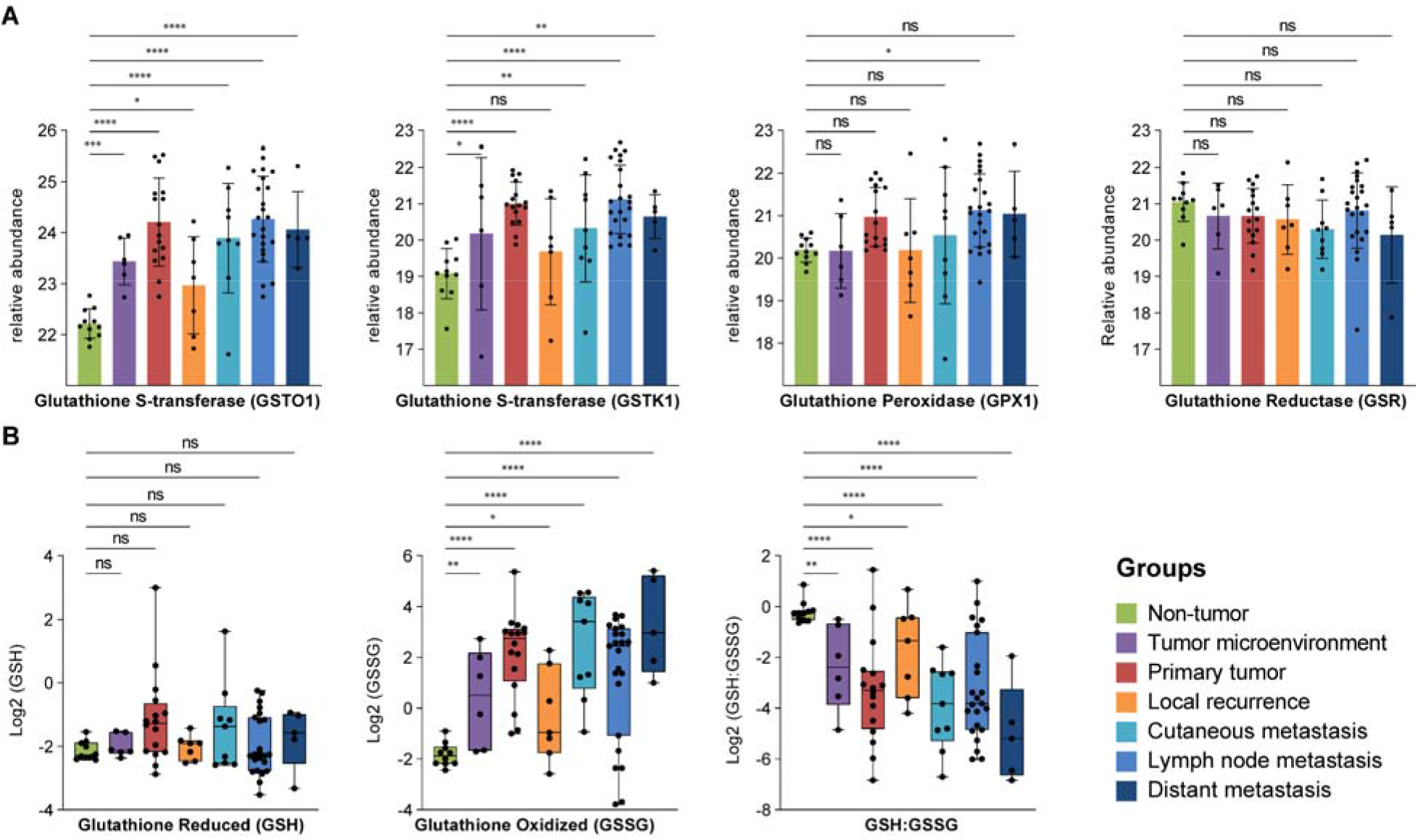
Melanomas and their microenvironment upregulate two members of the glutathione S transferase family GSTO1 and GSTK1 and the ratio between glutathione reduced and oxidized (GSH: GSSG) is downregulated in all groups of samples compared to non-tumors. *, **, ***, **** indicate significant differences compared to the non-tumor group (q-value < 0.05, 0.01, 0.001 and 0.0001 respectively), ns indicates not significant.

**Figure S7.**
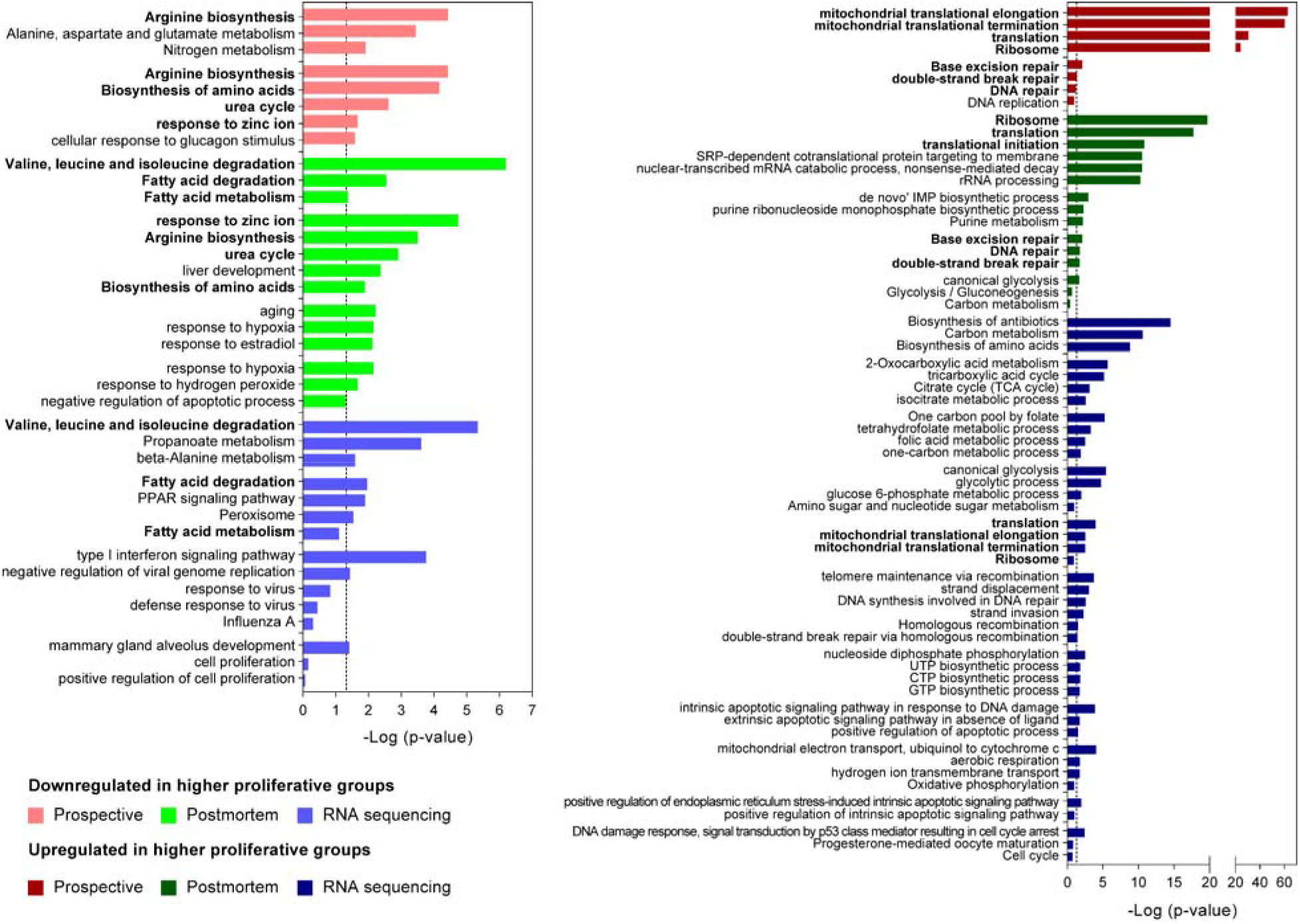
Significantly enriched biological processes and pathways in proteins dysregulated between the different proliferation-based groups in the three cohorts included in the study. The functional annotation enrichment analysis was performed using the online version of DAVID Bioinformatics V6.8.

**Figure S8.**
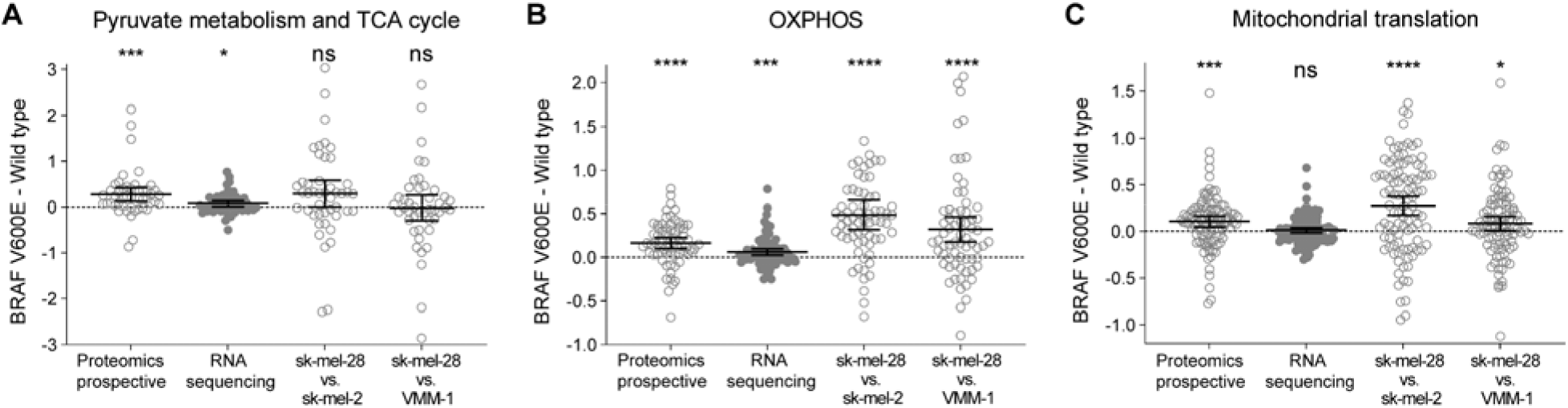
BRAF V600E mutation drives mitochondrial energy production via upregulation of OXPHOS. Difference plots and paired t-tests between BRAF V600E and wild type groups from the proteomic data of the prospective cohort and cell lines, plus the RNA sequencing data from the TCGA data. The groups of proteins analyzed belong to Pyruvate metabolism and TCA cycle, OXPHOS, and the mitochondrial translational machinery. Empty and filled circles correspond proteins and transcripts, respectively. The statistical significance of the paired t-test between BRAF status groups is represented (ns not significant, *, **, ***, and **** significantly different with p-value <0.05, <0.01, <0.001, and <0.0001 respectively).

## Notes

### Competing Interest Statement

The authors have declared no competing interest.

